# Characterization and structure of the human lysine-2-oxoglutarate reductase domain, a novel therapeutic target for treatment of glutaric aciduria type 1

**DOI:** 10.1101/2022.05.20.492856

**Authors:** João Leandro, Susmita Khamrui, Chalada Suebsuwong, Peng-Jen Chen, Cody Secor, Tetyana Dodatko, Chunli Yu, Roberto Sanchez, Robert J. DeVita, Sander M. Houten, Michael B. Lazarus

## Abstract

In humans, a single enzyme 2-aminoadipic semialdehyde synthase (AASS) catalyzes the initial two critical reactions in the lysine degradation pathway. This enzyme evolved to be a bifunctional enzyme with both lysine 2-oxoglutarate reductase (LOR) and saccharopine dehydrogenase domains (SDH). Moreover, AASS is a unique drug target for metabolic genetic diseases such as glutaric aciduria type 1 that arise from deficiencies downstream in the lysine degradation pathway. While work has been done to elucidate the SDH domain structurally and to develop inhibitors, neither has been done for the LOR domain. Here, we purify and characterize LOR, show that AASS is rate-limiting upon high lysine exposure of mice, and present the crystal structure of the human LOR domain, which should enable future efforts to identify inhibitors of this novel drug target.

## Introduction

The first step in lysine degradation via ε-deamination, also known as the saccharopine pathway, is catalyzed by 2-aminoadipic acid semialdehyde synthase (AASS). AASS is a bifunctional enzyme with an N-terminal lysine-oxoglutarate reductase (LOR) domain and a C-terminal saccharopine dehydrogenase (SDH) domain (**Figure 1A**), which are homologous to *Saccharomyces cerevisiae* LYS1 and LYS9, respectively.^1^ The LOR domain catalyzes the reductive deamination of L-lysine and 2-oxoglutarate into saccharopine (EC 1.5.1.8), which constitutes the first committed, and possibly rate-limiting, step in lysine degradation. SDH then oxidizes saccharopine into 2-aminoapidic semialdehyde and glutamate (EC 1.5.1.9). The bifunctional domain structure is conserved in all animals, but also *Dictyostelium discoideum* and plants, and seems to be associated with a function in lysine catabolism.^2^ Fungi such as *S. cerevisiae* use the saccharopine pathway for lysine biosynthesis.

**Figure 1.**
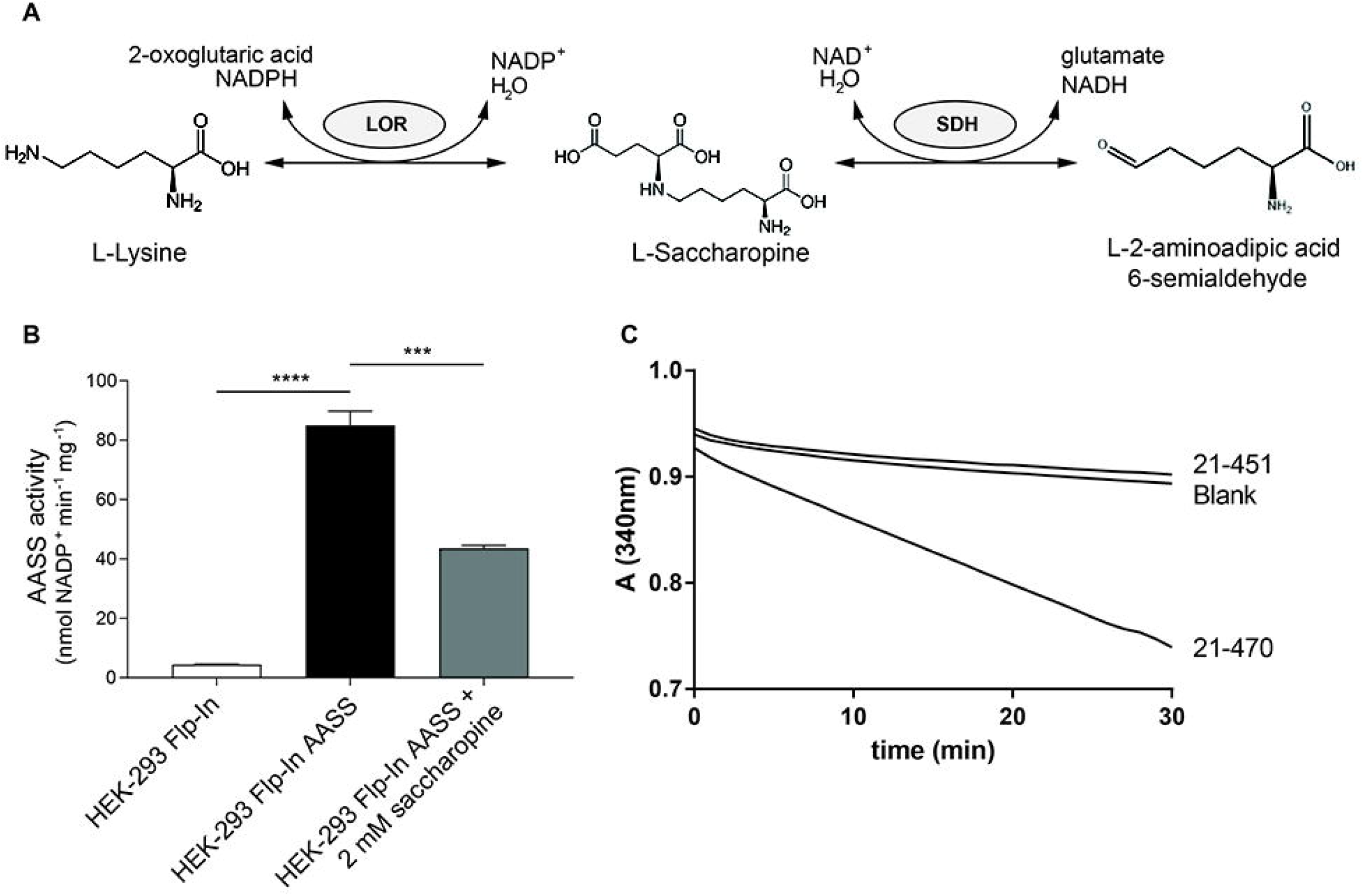
Characterization of AASS and the isolated LOR domain. (A) AASS reaction schema with the LOR and SDH activities. (B) Stable AASS expression in HEK-293 Flp-In cells and its inhibition by saccharopine. 10 mM 2-oxoglutarate (OG) was used as substrate. ***, P < 0.001; ****, P < 0.0001. Error bars indicate SD. (C) Progress curves showing catalytic activity of a short (amino acids 21-451) and long LOR construct (amino acids 21-470), with 1 mM OG, measured in triplicate. Only the average absorption at 340nm is displayed. The specific activity for 21-470 was 3.14 µmol/min.mg protein. Activity of the 21-451 construct was not detectable.

Several genetic inborn errors of metabolism occur in the lysine degradation pathway. Hyperlysinemia caused by AASS deficiency due to mutations in *AASS* (MIM 238700),^1,3,4^ is currently regarded as a biochemical phenotype of questionable clinical significance.^5^ This means that hyperlysinemia can be diagnosed through biochemical and genetic methods but is considered not harmful to the affected individual.^3,6-8^ In contrast, two other inborn errors of lysine degradation, glutaric aciduria type 1 (GA1) caused by mutations in *GCDH* (MIM 231670) and pyridoxine-dependent epilepsy caused by mutations in *ALDH7A1* (PDE-ALDH7A1; MIM 266100), are serious diseases. These diseases are caused by toxicity of the accumulating substrate of the defective glutaryl-CoA dehydrogenase (GCDH) and antiquitin, respectively. Dietary intervention to decrease lysine intake is one part of the treatment for both disorders, which reduces lysine degradation pathway flux. Given that current treatment for these diseases is not optimal and that there is also an endogenous (non-dietary) source of lysine (i.e. through protein degradation), we and others have hypothesized that GA1 and PDE-ALDH7A1 can be treated by pharmacological substrate reduction therapy, through inhibition of AASS.^9-11^ Although there is some debate on the contribution of AASS to the production of GCDH substrates in the brain ^10-14^, we previously showed that genetic inhibition of AASS, in cell lines and animal disease models of GA1, was highly effective in limiting the accumulation of GCDH substrates.^11^ Although brain glutarate remained higher than in WT mice, the level in *Gcdh*/*Aass* double KO mice was 4-fold lower when compared to *Gcdh* single KO mice.^11^ Since this therapeutic effect is comparable to the current treatment options, we believe AASS inhibition is a viable treatment strategy. Inhibition of the LOR domain is preferred given that it does not lead to accumulation of potentially toxic saccharopine.^15,16^

Given the interest in AASS as a new therapeutic target for lysine metabolic disorders such as GA1, there is great need for small molecule inhibitors that provide pharmacological proof-of-concept for efficacy of this approach. However, two major bottlenecks to drug discovery for this project remained: lack of a recombinant purification system; and a high-resolution crystal structure of the enzyme, which would enable high throughput and virtual screening, respectively. Structures have been reported for the isolated human SDH domain (e.g. PDB code 5O1O) and for the *S. cerevisiae* LYS1, which has ∼22% identity with the human LOR domain.^17-19^ However, the low sequence identity diminishes its usefulness for a drug discovery. Here, we describe enzyme and crystallographic studies including the first structure of the human LOR domain. We expect these results will enable the future development of a high affinity LOR inhibitor.

## Results

Pilot experiments aimed to overexpress AASS or its individual LOR and SDH domains in Escherichia coli indicated challenges in obtaining appreciable amounts of soluble protein. In order to characterize the protein, we initially transiently transfected HEK-293 cells with a plasmid encoding AASS with a C-terminal Myc-DDK tag (AASS-Myc-DDK) and used cell lysates as an enzyme source. Later, we generated and used Flp-In-293 cells stably overexpressing AASS-Myc-DDK. In both overexpression methods, LOR activity increased ∼30-fold when compared to control lysates and was reliably determined using a spectrophotometric enzyme assay (**Figure 1B**). LOR activity measured in the forward direction was inhibited by saccharopine consistent with its ability to compete with lysine and 2-oxoglutarate.^20^

The apparent steady-state kinetic properties of LOR in the full-length AASS-Myc-DDK were compared in the forward and reverse direction (**Table 1, Figure S1**). The rate of the reverse reaction was considerably slower than that of the forward reaction, which is consistent with previous reports.^21^ Positive cooperativity for several substrates is consistent with the reported tetrameric composition.^21,22^

**Table 1.** Steady-state kinetic properties of AASS-Myc-DDK and the LOR domain in forward and reverse reaction direction.

|  | Purified human liver | AASS-Myc-DDK |  |  | Isolated LOR |  |  |
| --- | --- | --- | --- | --- | --- | --- | --- |
| *Forward direction* | $K_{m, app}$<br>(mM) | $V_{max, app}$<br>(nmol min<sup>-1</sup> mg<sup>-1</sup>) | $K_{m, app}$<br>(mM) | n | $V_{max}$<br>(μmol min<sup>-1</sup> mg<sup>-1</sup>) | $K_m$<br>(mM) | n |
| L-lysine | 1.5 | 65 ± 2 | 11 ± 1 | 1.5 ± 0.1 | 28.8 ± 1.0 | 24 ± 2 | - |
| NADPH | 1 | 273 ± 99 | 0.42 ± 0.02 | 1.8 ± 0.1 | 65.4 ± 2.7 | 0.39 ± 0.02 | 2.1 ± 0.2 |
| 2-oxoglutarate | 0.08 | 106 ± 5<sup>a</sup> | 1.2 ± 0.1 | 1.5 ± 0.1 | 57.2 ± 1.0 | 1.4 ± 0.1 | - |
| *Reverse direction* | $K_{m, app}$<br>(mM) | $V_{max, app}$<br>(nmol min<sup>-1</sup> mg<sup>-1</sup>) | $K_{m, app}$<br>(mM) | n | $V_{max}$<br>(nmol min<sup>-1</sup> mg<sup>-1</sup>) | $K_m$<br>(mM) | n |
| saccharopine | 1.5 | 7.7 ± 0.1 | 1.0 ± 0.1 | 1.9 ± 0.1 | 443 ± 6 | 1.7 ± 0.1 | 2.2 ± 0.1 |
| NADP<sup>+</sup> | NR | 7.4 ± 0.4 | 1.0 ± 0.1 | - | 525 ± 5 | 0.42 ± 0.01 | 1.6 ± 0.1 |

The $K_{m, app}$ of partially purified AASS from human liver has been reported<sup>20,34</sup>. The pH optima in these studies were 7.8 for LOR forward and between pH 8.8-9.5 for LOR reverse. n is the Hill coefficient. <sup>a</sup>Substrate inhibition. Values represent mean ± SD.

In order to optimize production of recombinant isolated LOR protein in *E. coli*, we evaluated several versions of His-tagged proteins for solubility and activity. We screened different length constructs in order to determine the appropriate domain boundaries. We were able to obtain large amounts of active soluble protein using an N-terminal His-sumo-tag construct with the mitochondrial transit peptide membrane segment removed (amino acids 21-470, **Figure 1C**). This recombinant LOR was subsequently used to determine the steady-state kinetic properties. Overall, the kinetic constants resembled those of full-length AASS-Myc-DDK with the forward reaction velocity being considerably higher than the reverse (**Figure S1, Table 1**).

If an enzyme is truly rate-limiting in a pathway, the concentration of the enzyme (i.e. expression level) determines flux. This can be evaluated by titrating the activity of an enzyme with a specific, irreversible inhibitor.^23^ Such an inhibitor is currently not available for AASS. In order to start addressing the question whether AASS is rate-limiting in vivo, we used the *Aass* KO mouse model.^11^ If AASS is a rate-limiting enzyme in vivo, *Aass*^+/-^ mice, which have a 50% reduction in AASS expression, should have decreased lysine degradation flux. We have previously measured key metabolites in a cohort of animals carrying *Gcdh* and *Aass* KO alleles on a chow diet.^11^ We now grouped all mice according to their *Aass* genotype. As expected, *Aass* KO mice had elevated plasma lysine concentrations (**Figure 2**). Plasma lysine in *Aass*^+/-^ mice was comparable to plasma lysine in *Aass*^+/+^ mice. This result argues against AASS catalyzing the rate-limiting step in lysine catabolism under standard chow fed conditions. However, we then exposed these mouse models to high lysine through diet and drinking water. Under these conditions, the number of *Aass* KO alleles explains a significant percentage (50%) of the variation in plasma lysine concentrations with *Aass*^+/-^ animals clearly higher than those of *Aass*^+/+^ animals. Combined, these data suggest that AASS can be rate-limiting under conditions of high lysine load.

**Figure 2.**
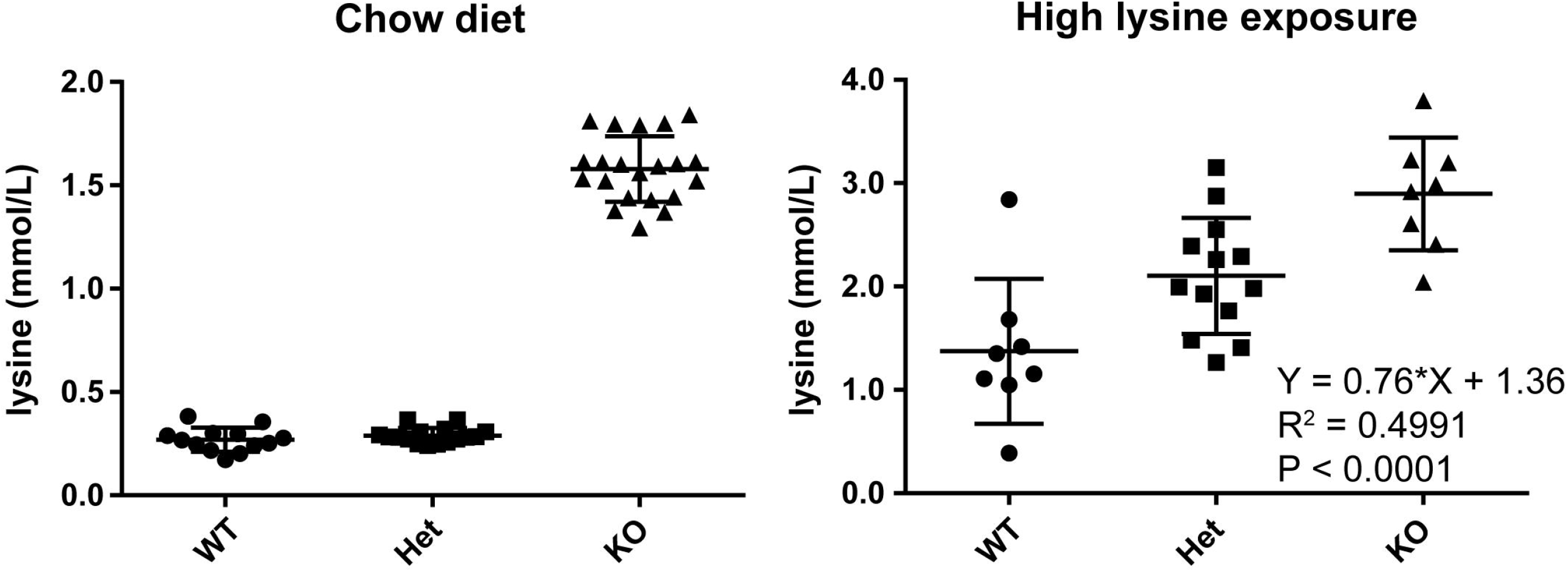
AASS is rate limiting upon high lysine exposure in mice. Plasma lysine concentration in mice on chow diet and upon high lysine exposure. Mice are grouped according to their genotype at the *Aass* locus (*Aass*^+/+^, *Aass*^+/-^ and Aass-/-). Each group contains *Gcdh*^+/+^, *Gcdh*^+/-^ and *Gcdh*^−/-^ animals. Additional statistical analysis through a 2way ANOVA showed that the *Gcdh* locus did not explain variation in plasma lysine concentration (Interaction: 4.79%, not significant; *Gcdh* genotype: 0.15%, not significant; and *Aass* genotype: 52.05%, P = 0.0001). A Tukey’s multiple comparisons test shows significant differences between WT, Het and KO group (WT vs Het, P<0.05; WT vs KO, P<0.001; and Het vs KO, P<0.05).

No human structures have been reported for AASS/LOR, and the sequence identity with S. cerevisiae LYS1 is relatively low making structural prediction for the human enzyme challenging. We initially purified a short construct (amino acids 21-451) that expressed well in E. coli with high purity and a monodisperse peak on gel filtration. We screened conditions and obtained crystals that after optimization diffracted to 2.2 Å and solved the structure (PDB XXXX) by molecular replacement using the yeast LYS1 structure as a search model (**Table S1**). The overall structure of the human protein is quite similar to yeast LYS1 despite the low sequence identity, indicating strong conservation of function despite the evolved differences such as adopting bifunctional enzyme architecture and switching from NAD^+^ to NADP^+^. As seen in **Figure 3A**, the LOR domain itself consists of two lobes, both Rossman folds with a central beta sheet with alpha helices on the outside. The lobes are connected twice with the C-terminus ending up in the N-terminal lobe. Some of the extended loops and predicted catalytic residues were not visible in this structure. Moreover, when we tested this short construct for activity, we discovered that, despite the fact that it was well-folded and aligned nicely with the yeast structure, it had no activity in our in vitro assay (**Figure 1C**).

**Figure 3.**
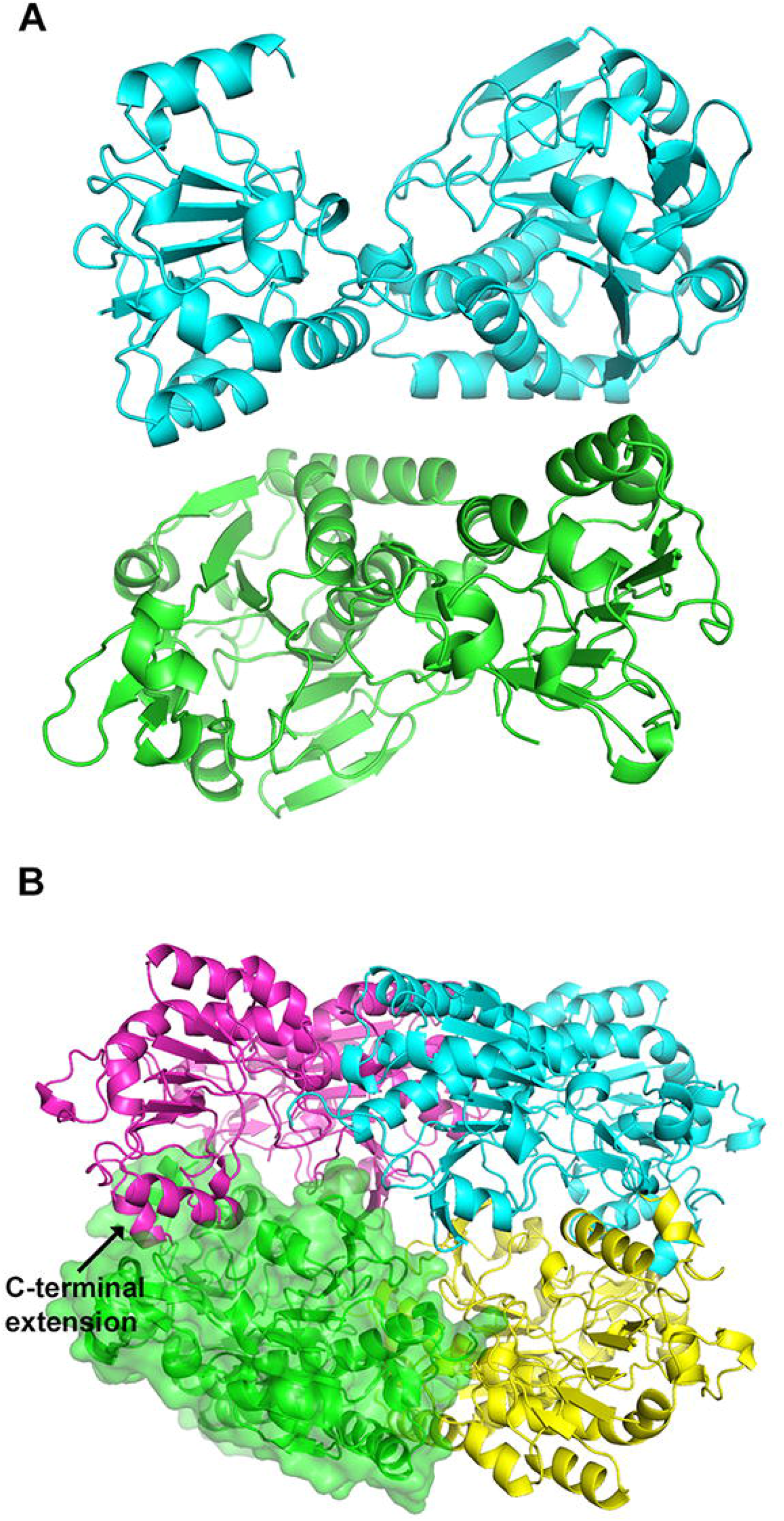
Overall structure of two LOR constructs. (A) Asymmetric unit of the short inactive construct shows a dimer of the LOR domain of AASS. The symmetry mate dimerization interface (**Figure S3**) is likely physiological. (B) Tetrameric structure of the long LOR constructs shown the four monomers assembling into a compact sphere. A surface is shown for one of the monomers to highlight the packing.

Therefore, to obtain an LOR structure of a protein that maintained catalytic activity, we focused on the longer construct (amino acids 21-470). This initial construct crystallized readily but did not diffract well, therefore we introduced five surface mutations to induce better packing. We were then able to obtain a crystal structure of the longer active construct, solving the structure (PDB YYYY) by molecular replacement with the short construct. The longer construct crystallized in a new space group as a tetramer (**Figure 3B**). Each monomer is very similar to the short construct with a root-mean-square deviation (RMSD) of 0.5691 Å^2^ between the most similar chains.

Several lines of evidence support an oligomeric state for AASS. Both constructs eluted on gel filtration on a Superdex200 Increase column at around 12.5 mLs, which equates to an approximate molecular weight between 150-200 kDa (**Figure S2**). This would suggest either a trimer or tetramer. Secondly, the dimeric interface is quite large. According to the PISA server, the average dimer interface of the longer construct is 1810 Å^2^ with a predicted ΔG of -6 kcal/mol. We reanalyzed the structure of the short construct and noticed that although the interface in the asymmetric unit is likely non-physiologic, the interface of a monomer with its crystallographic symmetry-mate is identical to the long interface in the tetramer (**Figure S3**), further supporting the validity of this interface. Next, the kinetic analysis of the recombinant LOR enzyme is also consistent with a multimer, given the cooperativity observed for NADP^+^ (**Figure S1**).

At a structural level, the C-terminus of the longer construct forms an alpha helix that penetrates into the active site of the adjacent monomer. One paradox to resolve was why the C-terminus is so critical for activity, despite the facts that it is distant from the active site and the shorter construct is well folded. One hypothesis is that the C-terminus interacts with the catalytic residues of an adjacent monomer with the enzyme active only in an oligomeric state. Early on, it was reported in the literature that full-length human AASS exists as a tetramer when isolated from liver,^24^ consistent with our observations. Now, we have structural information on the tetrameric interface.

Compared to the yeast structure, the human one has two large insertions, both in the C-terminal lobe. First, there is a 37-amino acid insertion after residue 262. We see good density for this insertion, which forms two additional short alpha helices that extend away from the tetramer. The second is an insertion after residue 375, which forms a beta sheet extension in the C-terminal lobe. In the structure of the long construct, we see electron density at the interface of these loops from two different monomers that sit between Asp 388 sidechains from each monomer and Glu 373 carbonyl backbones from each monomer (**Figure S4**). We have assigned this density to a magnesium ion which could help neutralize the negative charge from the aspartates. It is most likely hydrated given the distance between the aspartates, but we cannot see enough density for the water shell at the resolution of this structure.

Another important difference between LYS1 and LOR is that the human protein evolved to use NADPH while yeast uses NADH. This allows the LOR to drive the reaction toward production of saccharopine due to the much higher (essentially fully reduced) ratio of NADPH/NADP^+^ compared to NADH/NAD^+^. Therefore, we would expect changes in the binding pocket to accommodate the phosphate group on NADP^+^. We were unable to obtain a crystal structure with substrates bound, presumable due to the low affinity to AASS for all of its substrates. However, the structure overlays favorably with the yeast structure bound to NAD^+^ and therefore we can predict the key interactions. Initial alignments^18^ predicted the human protein has a glutamate where the yeast binds the ribose of NADH. However, in the structure we can see that the Ser 266 and Arg 267 sidechains match up to where the aspartate resides in the yeast structure (**Figure S4**). This explains why the human protein can bind the NADPH, replacing the negative charge of the yeast loop with a hydroxyl group (of serine 266) and a positively charged arginine 267 to interact with the additional phosphate of NADPH.

Hyperlysinemia in humans has been associated with the p.R65Q, p.A154T, and p.L419R variants in the LOR domain of AASS.^3^ Our earlier work showed that many of these mutations lead to loss of protein expression,^3^ and with our structure we can better understand the consequences of these amino acid substitutions on the LOR structure. Arg 65 makes several important contacts including a salt bridge to Asp 69 and a hydrogen bonding interaction with the carbonyl backbone of Gln 60, helping to stabilize several portions of the N-lobe. Ala 154 is located exactly where the nicotinamide moiety of NAD^+^ binds in the yeast structure, so the threonine would likely preclude its binding. The Alanine-Glycine motif from 154-155 is conserved from the yeast protein to the human one, likely because any larger sidechains would obstruct nicotinamide binding. Lastly, Leu 419 is buried in a hydrophobic pocket in the N-lobe, so an arginine sidechain would likely destabilize the pocket.

Since AASS is a viable therapeutic target for GA1, our next objective was to develop and validate an assay amenable to high throughput screening for identifying inhibitors of the LOR domain. We used the absorption at 340 nm to monitor NADPH consumption in a 96-well plate format. In order to identify compounds to help validate the assay, we performed a virtual screen to identify potential active site inhibitors to assess viability of the assay. We purchased 126 commercially available potential hit compounds identified by virtual screen, then tested them in the LOR enzyme assay. As seen in **Figure 4A**, almost all compounds had minimal effect on activity. However, we identified one compound, **105**, that showed inhibition several standard deviations away from the average. We then re-synthesized racemic compound 105 which contains one chiral center (**Figure 4B**) to confirm its activity. We also prepared both 105 enantiomers found that the *S*-105 enantiomer was more active against both the recombinant LOR protein and the full-length AASS construct. Additional compounds synthesized did not show improved potency (**Table S2** and **Figures S5-S7**). However, while the potency of this compound was modest, it demonstrated that the assay was robust, dose responsive and useful to identify inhibitors. There is also the possibility to further miniaturize to a 384-well plate format with incubation at room temperature for high-throughput screens. We expect these results to enable discovery of more potent inhibitors of LOR using an *unbiased* large library screen for identification of both active or allosteric site inhibitors.

**Figure 4.**
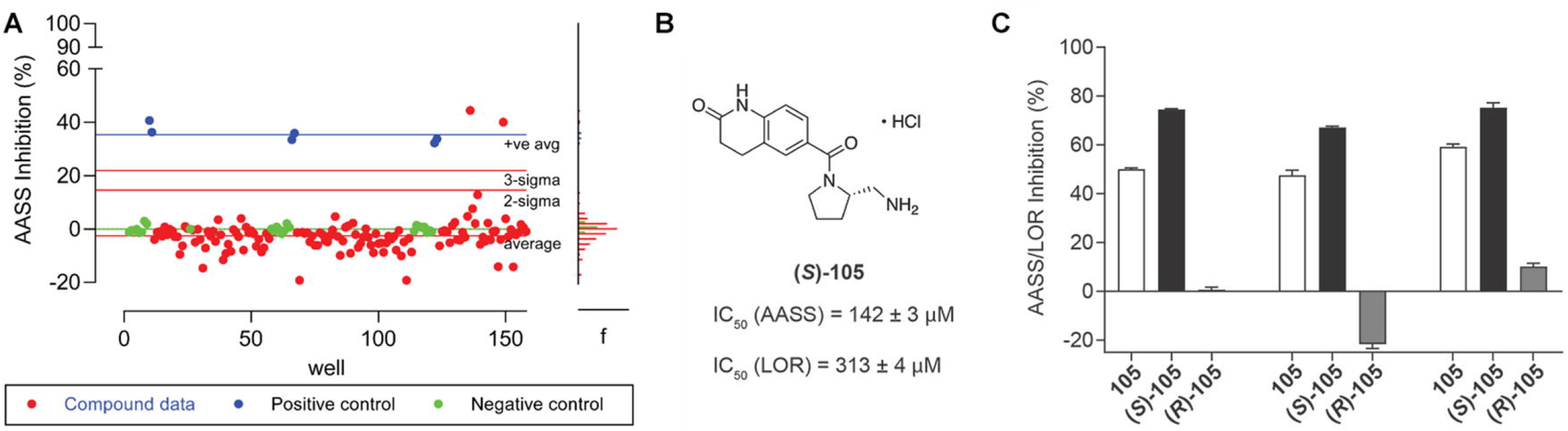
Development of LOR assay to screen for inhibitors. A) Signal to noise of assay shows good properties for a screen, with standard deviation lines noted in red lines. The Z-factor is typically between 0.7 and 0.8. B) Structure and IC_50_ of AASS/LOR inhibitor. (C) Full-length AASS inhibition in the forward reaction and LOR domain inhibition in the forward and reverse reaction by inhibitor 105 and its stereoisomers at a concentration of 300 µM. 1 mM 2-oxoglutarate was used as a substrate.

## Discussion

AASS is an important enzyme in lysine degradation. It catalyzes the first committed step in this pathway and is a potentially attractive pharmacological target for the treatment of two inborn errors of metabolism, GA1 and PDE-ALDH7A1. Herein, we developed chemical and biological tools that can be useful for the future development of a high affinity inhibitor of the LOR domain of AASS. We overexpressed AASS-Myc-DDK, purified a recombinant isolated LOR domain, and determined the kinetic constants of these enzymes. In a virtual screen, using a homology model of LOR, we identified one molecule that is able to inhibit enzyme activity with modest inhibition activity (IC_50_ 142 µM) and then established preliminary structure-activity relationships. We solved the first crystal structure of the human LOR domain at 2.2Å resolution.

AASS catalyzes the first committed, and possibly rate-limiting, step in lysine degradation. Although the LOR reaction is reversible in vitro, in vivo it is likely driven toward the production of saccharopine by the nearly fully reduced NADPH/NADP^+^ ratio in mitochondria. A defect in the mitochondrial biosynthesis of NADP^+^ leads to hyperlysinemia demonstrating that in man and mouse NADPH cannot be replaced by NADH.^25-28^ There is evidence that limited 2-oxoglutarate availability in some inborn errors of metabolism, such as urea cycle defects, limits lysine degradation,^29^ but it is unknown if 2-oxoglutarate is limiting under non-pathological conditions. The affinity of AASS for lysine is relatively low. The previously reported K_m_ of partially purified AASS/LOR from human liver was 1.5 mM ^20^ (Table 1). We calculated a K_m_ of 11 mM for AASS-Myc-DDK and 24 mM for isolated LOR. Plasma concentrations of lysine are usually lower than 300 µmol/L^3,11^. Estimated tissue concentrations for lysine are very similar (336 µmol/L for liver and 142 µmol/L for heart based on Reference^30^). The low affinity of AASS/LOR for lysine in combination with the estimated tissue lysine concentration suggests that the velocity of lysine degradation will show a linear relationship to the mitochondrial lysine concentration. However, a gene-dose effect on plasma lysine concentration in mice carrying one *Aass* null allele was only evident upon high lysine exposure. No such effect was observed in mice on chow diet, which likely supplies lysine at quantities close to the actual nutritional requirement. Therefore, we speculate that with a relatively low, but adequate lysine supply, the rate of its degradation is determined by cellular lysine uptake and subsequent protein synthesis and not by AASS activity. Although there is little available information on lysine transport across the plasma and mitochondrial membrane, it is likely driven by lysine concentration in a saturable process.^31,32^ In contrast, under conditions with a larger supply of lysine, for example during catabolism or postprandial, AASS/LOR catalyzes the rate-limiting step in lysine degradation and thus reinforces the notion that inhibiting of LOR is a suitable strategy in the treatment of GA1. Indeed, it has been reported that during acute illness the urinary excretion of glutaric and 3-hydroxyglutaric acid can rise dramatically with values 4-fold higher than observed as compared to baseline illness. ^33^

To conclude, we have characterized and purified the LOR domain of the important metabolic enzyme AASS. We show that AASS can be rate-limiting in the pathway and present the first crystal structure of the human LOR domain. We established an assay to measure inhibition of AASS in high throughput and identified a weak inhibitor by virtual screen that helped validate our assay. The development of a recombinant purification system and a high-resolution crystal structure and will enable future efforts to further validate this enzyme as a potential therapeutic target for the treatment of GA1 and will enable improved inhibitor discovery to provide pharmacological proof of concept for efficacy in cells and in vivo.

## Supporting Information

A complete description of the methods and supplementary figures and tables are in Supporting Information.

## AUTHOR INFORMATION

### Conflict of Interest

The authors declare no conflict of interest.

## ACKNOWLEDGMENT

Research reported in this publication was supported by the Eunice Kennedy Shriver National Institute Of Child Health & Human Development and the National Institute Of General Medical Sciences of the National Institutes of Health under Award Numbers R21HD102745 (to S.M.H. and R.J.D.) and R35GM124838 (to M.B.L.). The content is solely the responsibility of the authors and does not necessarily represent the official views of the National Institutes of Health. This research used resources of the National Synchrotron Light Source II, a U.S. Department of Energy (DOE) Office of Science User Facility operated for the DOE Office of Science by Brookhaven National Laboratory, under Contract No. DESC0012704. The Life Science Biomedical Technology Research resource is primarily supported by the NIH, National Institute of General Medical Sciences (NIGMS) through a Biomedical Technology Research Resource P41 grant (No. P41GM111244), and by the DOE Office of Biological and Environmental Research (No. KP1605010). This work utilized the NMR Spectrometer Systems at Mount Sinai acquired with funding from National Institutes of Health SIG grants 1S10OD025132 and 1S10OD028504.

## Notes

### Competing Interest Statement

The authors have declared no competing interest.

