## Supplemental methods, supplemental figure and supplemental table 1 for "Characterization and structure of the human lysine-2-oxoglutarate reductase domain, a novel therapeutic target for treatment of glutaric aciduria type 1"

Table of contents

| | |
|---|---|
| Supplementary figures and legends | Pages 2-11 |
| Supplementary tables | Pages 12-13 |
| Materials and Methods | Pages 14-18 |
| Synthesis of AASS inhibitors | Pages 19-31 |
| NMR spectra | Pages 32-48 |
| References | Page 49 |

**Supplementary Figures**

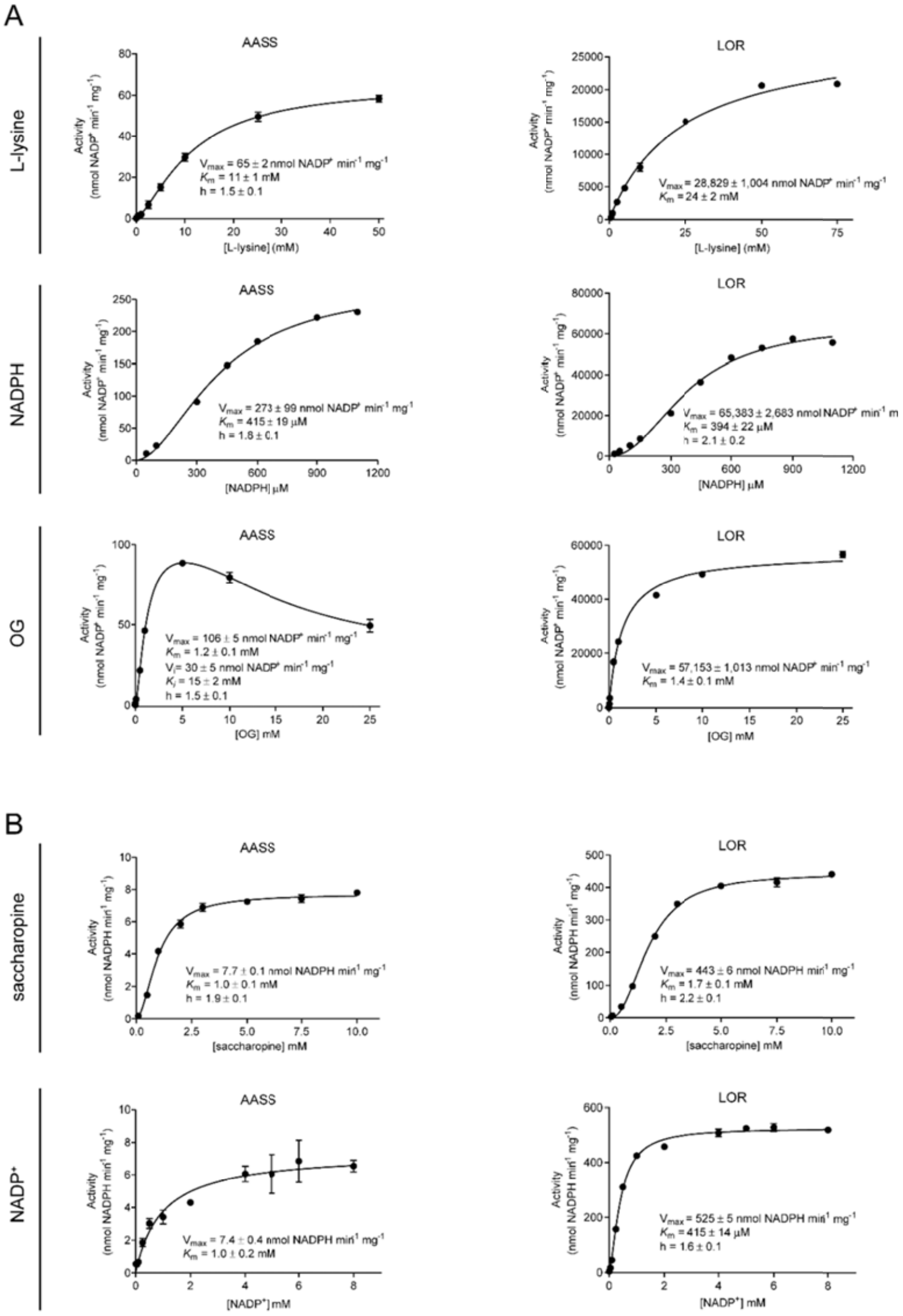

**Figure S1.** Enzyme kinetics of human AASS and LOR. (A) The effect of substrates L-lysine, NADPH and 2-oxoglutarate (OG) concentration on the catalytic activity of the forward reaction of AASS and LOR. The AASS/LOR activity was assayed at standard conditions (0 – 50 or 15 mM L-lysine, 0 – 1200 or 300 μM NADPH, 0 – 25 mM or 10 mM OG and 37 °C). (B) The effect of substrates saccharopine and $NADP^+$ concentration on the catalytic activity of the reverse reaction of AASS and LOR. The AASS/LOR activity was assayed at standard conditions (0 – 10 or 2 mM saccharopine, 0 – 8 mM or 1 mM $NADP^+$ and 37 °C). The data were analyzed by non-linear regression analysis using the GraphPad

Prism 7 software and the Michaelis-Menten equation or Hill equation. The effect of OG in AASS was analyzed using the modified Hill equation of LiCata and Allewell to account for substrate inhibition [1]. The mean ± SD values are shown (see also Table 1).

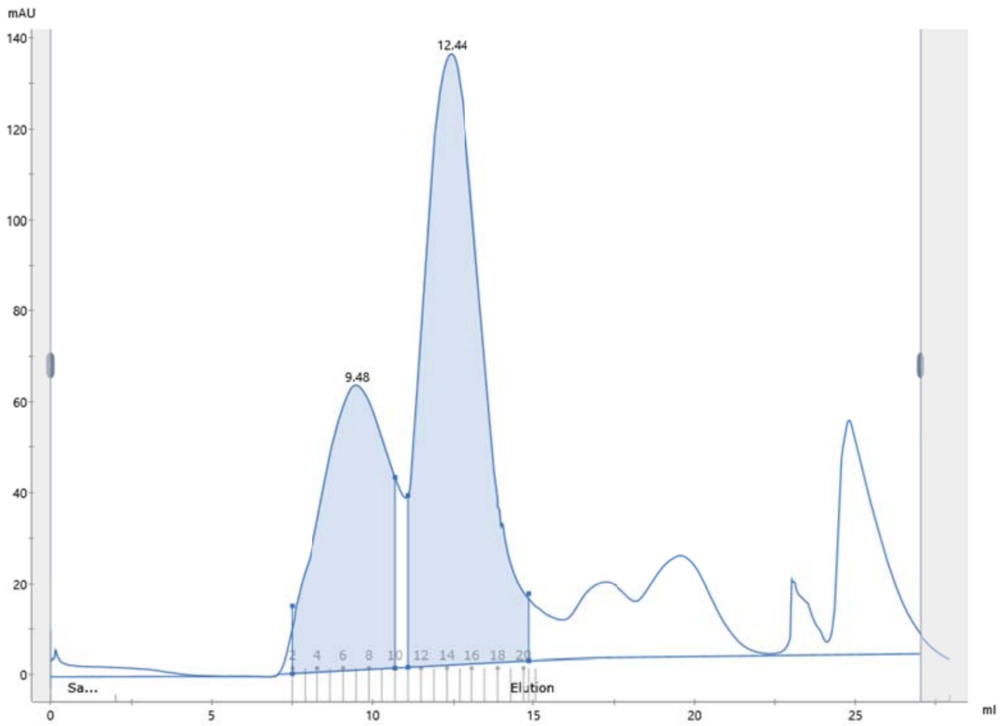

**Figure S2. FPLC chromatograph of LOR domain.** The protein was purified on a Superdex200 size exclusion column after removal of the His-sumo tag. The retention time of 12.44 ml on the Superdex200 10/300 GL column is consistent with a molecular weight of approximately 150-200 kDa. This suggests the LOR domain itself (50 kDa) forms an oligomer in solution.

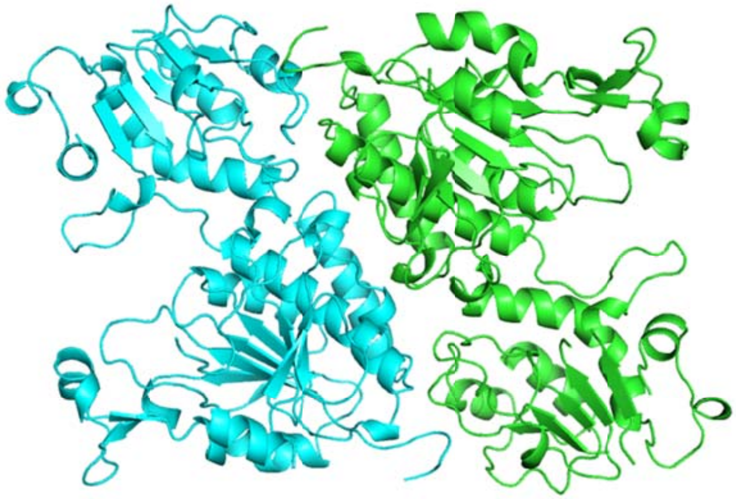

**Figure S3. Symmetry interface of short construct.** We analyzed the interface of a monomer in the short construct with its symmetry-mate and observed a nearly identical alignment to the tetramer in the long construct, with greater than 1800 $Å^2$ of interface area.

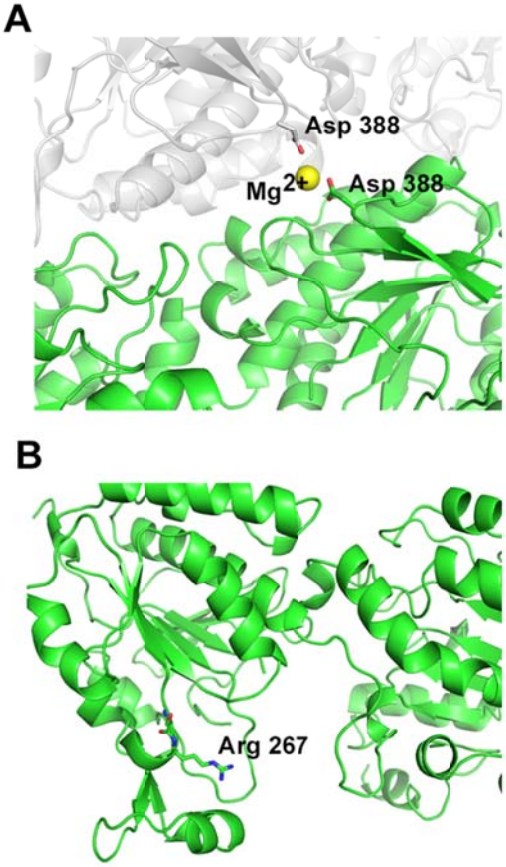

**Figure S4. Functional details of the structure**. A. Dimerization interface with a likely divalent cation mediating two Asp 388 residues. This likely contributes to the energy of tetramerization. B. Likely NADPH binding site, with Arg 267 shown. This residue is not conserved in yeast which binds NADH.

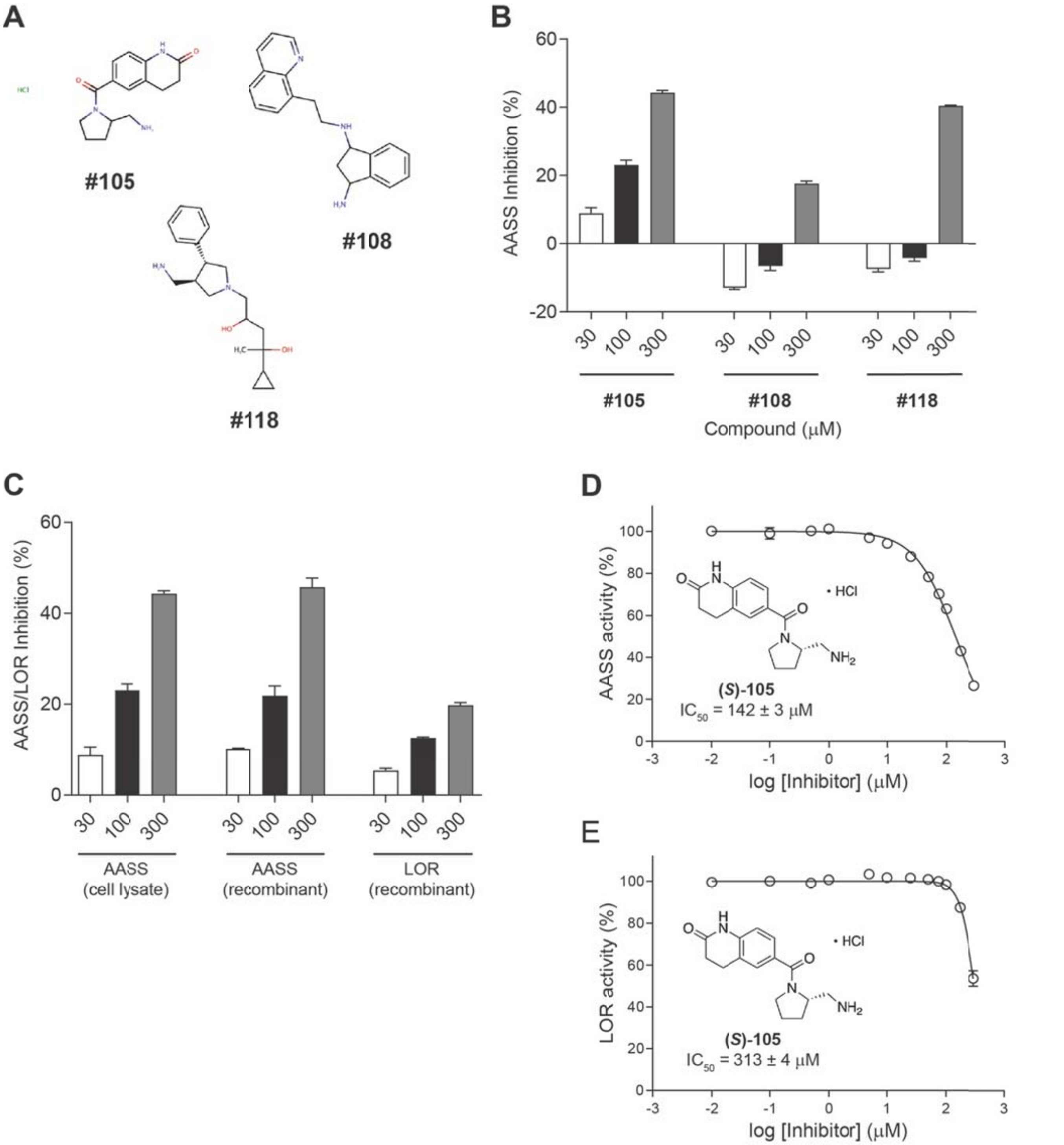

**Figure S5.** AASS/LOR inhibition. (A) Chemical structures of the 3 out of 126 virtual screening hit compounds that exhibited AASS inhibition above 10%. (B) Dose-response inhibition of AASS by the 3 virtual screening hit compounds tested at 30, 100 and 300 μM. (C) Confirmation of AASS/LOR inhibition by hit compound #105 using full-length AASS from cell lysate, recombinant full-length AASS and recombinant LOR. (D and E) Chemical structure and dose-response curves of compound #(*S*)-105 stereoisomer upon inhibition of AASS from cell lysate (D) and recombinant LOR (E). Compound #105 and #(*S*)-105 stereoisomer for this experiment were synthesized in-house.

**A** **SAR by catalog (Table S1B)**

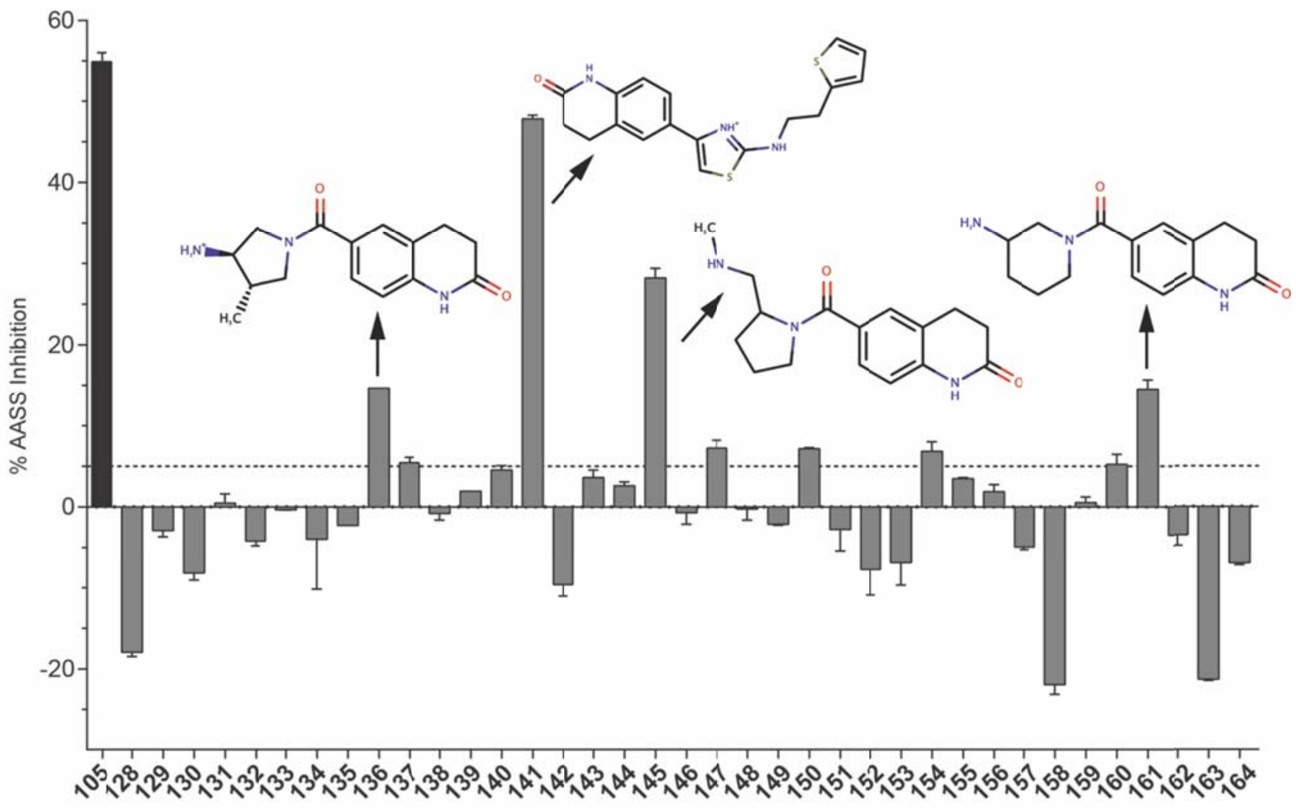

**B**

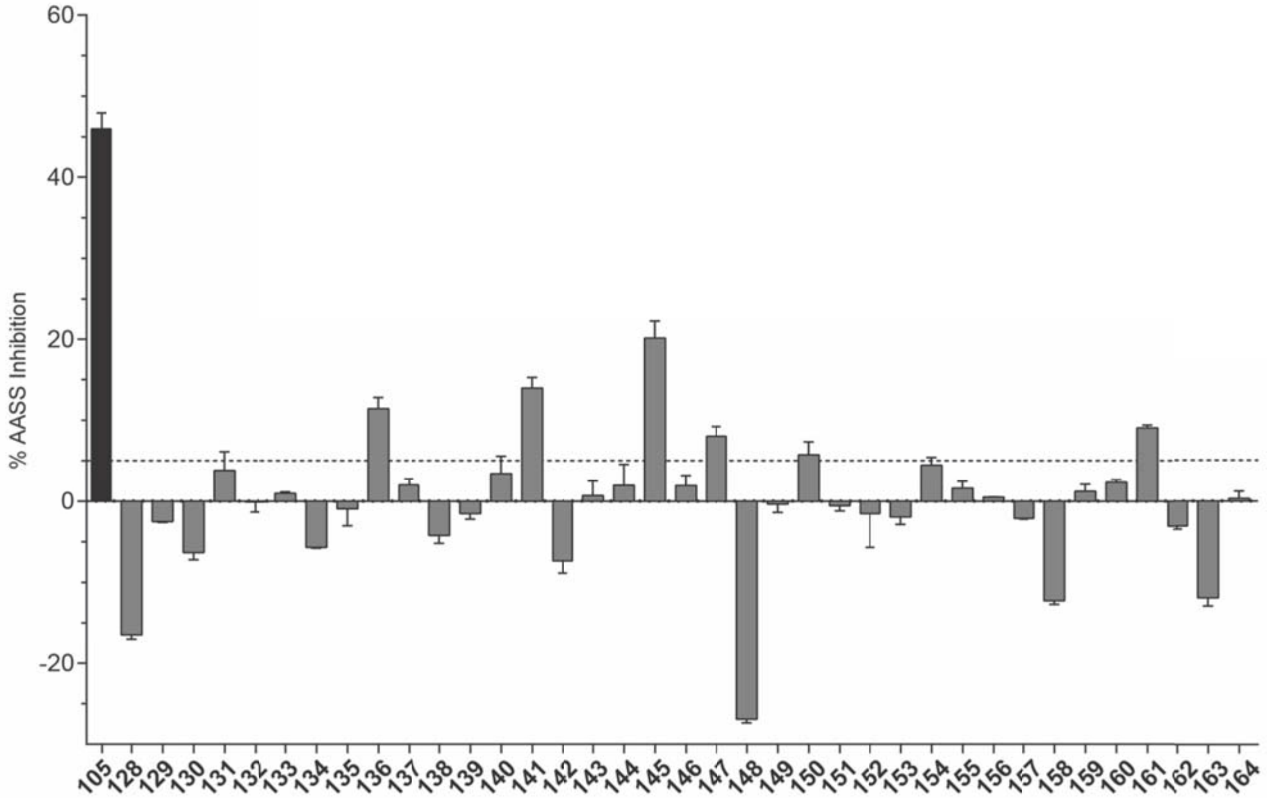

**Figure S6.** SAR by catalog for AASS/LOR inhibition. (A) Inhibition of LOR (in %) for 37 analogs selected based on similarity and/or potency. Compounds were tested at 300 μM and 1 mM OG was used as substrate. (B) Inhibition of LOR (in %) for 37 analogs selected based on similarity and/or potency. Compounds were tested at 300 μM and 10 mM OG was used as substrate. A cell lysate of HEK-293 Flp-In cells stably overexpressing AASS was used.

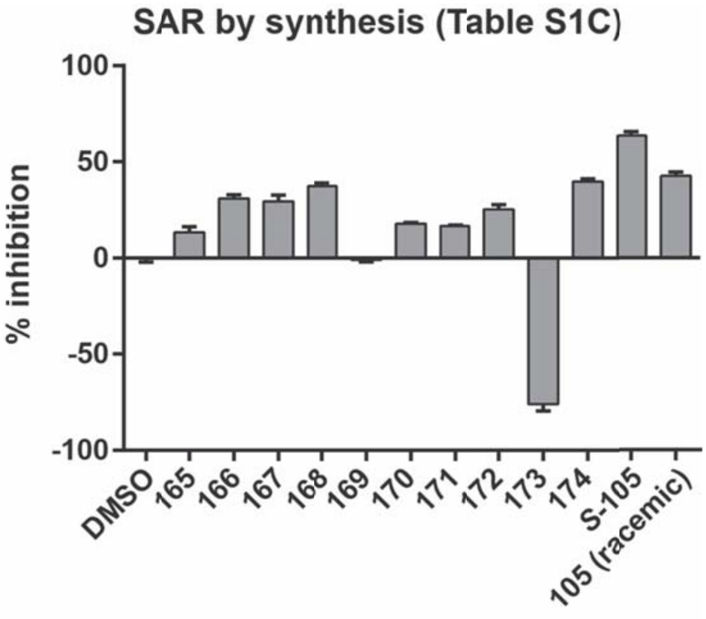

**Figure S7.** SAR through synthesis of selected compound #105 analogs. Inhibition of LOR (in %) for 10 analogs selected based on similarity. Compounds were tested at 300 μM and 1 mM OG was used as substrate with recombinant LOR as protein source.

#### Supplementary Tables.

#### Table S1. Data collection and refinement statistics

| | LOR short construct | LOR long construct |
|---|---|---|
| **Data collection** | | |
| Space group | P 21 21 2 | P 1 21 1 |
| Cell dimensions | | |
| *a*, *b*, *c* (Å) | 77.356, 154.715, 70.674 | 73.005, 131.944, 96.494 |
| α, β, γ (°) | 90, 90, 90 | 90, 100.723, 90 |
| Resolution (Å) | 42.91 - 2.18 (2.258 - 2.18)* | 47.4 - 2.65 (2.745 - 2.65) |
| $R_{sym}$ or $R_{merge}$ | 0.172 (1.089) | 0.407 (2.385) |
| $I / \sigma I$ | 4.4 (1.0) | 2.7 (0.8) |
| Completeness (%) | 96.7 (98.0) | 96.4 (97.0) |
| Redundancy | 3.1 (2.9) | 5.6 (5.6) |
| $CC_{1/2}$ | 0.981 (0.367) | 0.974 (0.431) |
| **Refinement** | | |
| Resolution (Å) | 42.918 - 2.18 (2.228 - 2.18) | 47.412 - 2.65 (2.701 - 2.65) |
| No. reflections | 43247 (4293) | 50070 (5086) |
| $R_{work}$ / $R_{free}$ | 0.2136 / 0.2450 | 0.2780 / 0.3192 |
| No. atoms | 6575 | 13532 |
| Protein | 6326 | 13490 |
| Ligand/ion | 0 | 2 |
| Water | 249 | 40 |
| *B*-factors | | |
| Protein | 43.89 | 46.25 |
| Ligand/ion | | 29.08 |
| Water | 38.18 | 20.57 |
| R.m.s. deviations | | |
| Bond lengths (Å) | 0.003 | 0.002 |
| Bond angles (°) | 0.57 | 0.52 |

*Values in parentheses are for highest-resolution shell. Short construct structure was from a single crystal. The long construct was from three crystals.

**Table S2. Summary of compounds tested against AASS**

Worksheet S2A. Compounds tested in the virtual screen.

Worksheet S2B. SAR by catalog.

Worksheet S2C. SAR by synthesis.

See attached Table S2 Virtual screening compounds and SAR.xlsx

#### Materials and Methods

*Materials*

Hepes, 2-oxoglutarate, NADPH, KOH, $NADP^+$, $NH_4Cl$, Tris base, DMSO anhydrous and saccharopine were from Millipore-Sigma. Triton X-100 was obtained from Fisher Scientific. HEK-293 cells were obtained from American Type Culture Collection (Manassas, VA, USA). Flp-In™-293 Cell Line, Flp-In vectors, zeocin and hygromycin B were obtained from Invitrogen (now Thermo Fisher Scientific). DMEM, penicillin, streptomycin and fetal bovine serum were also obtained from Thermo Fisher Scientific (Waltham, MA, USA). All chemicals were of analytical grade.

*Generation of a Flp-In-293 cell line stably overexpressing AASS*

Human *AASS* (NM_005763) with a C-terminal Myc-DDK tag in the pCMV6-Entry vector was purchased from Origene (RC224831) and used for transient transfection. The Flp-In Core System and Flp-In-293 cell line were obtained from Invitrogen (Thermo Fisher Scientific). The full length Myc–DDK tagged *AASS* cDNA insert was released from the pCMV6 vector with KpnI (New England Biolabs; NEB) and PmeI (NEB) and purified by QIAquick Gel Extraction kit (QIAGEN). The pcDNA5/FRT vector was linearized with BamHI (NEB), purified by Monarch DNA Gel extraction kit (NEB) and blunt ends were created using the Quick Blunting kit (NEB). The resulting vector was purified, digested with KpnI and again purified. The resulting AASS insert and pcDNA5/FRT vector were ligated by using the Quick Ligation Kit (NEB) and transformed to CopyCutter EPI400 chemically competent cells (Lucigen). Sequencing revealed that the resulting construct had lost AA of the ochre stop codon. This was corrected by using Q5 Site-directed Mutagenesis Kit (NEB) with primers: Fw: 5'-aaTCCACTAGTCCAGTGTGG-3' and Rev: 5'-AAACCTTATCGTCGTCATC-3'. The AASS-pcDNA5/FRET was transformed to CopyCutter EPI400 chemically competent cells and the sequence was verified by sequencing. Both the AASS-pCDNA5/FRT and pOG44 plasmid DNA were purified by NucleoBond Xtra Midi Plus EF DNA purification Kit (Macherey-Nagel).

Purified pOG44 and AASS-pcDNA5/FRT plasmids were co-transfected into Flp-in-293 cells with a 1:9 ratio by using Lipofectamine 2000 reagent (Invitrogen, Thermo Fisher Scienific). Flp-in-293 cells were cultured in DMEM media supplemented with 10% FBS, Pen/Strep and 100 μg/mL Zeocin. At the day of transfection, incubation media were removed, cell were washed with warm PBS and then DMEM media supplemented with 10% FBS, Pen/Strep, but without Zeocin were added. Transfected cells were incubated at 37°C with 5% $CO_2$ for 24h and transfection media were then replaced by fresh DMEM media supplemented with 10% FBS and Pen/Strep and cells were incubated for next 24h. After 48h, cells were split to 25% confluence into new Hygromycin B selective media: DMEM supplemented with 10% FBS, Pen/Strep and 100 (200) μg/mL Hygromycin B. Cells were incubated for 2-3 weeks at 37°C with 5% $CO_2$. Selective media were replaced every 3-4 days. After Hygromycin B resistant foci were formed, cells were transferred to new flasks and were grown on Hygromycin B selective media until they were 75-80% confluent. Cell pellets were collected to confirm overexpression of AASS using immunoblotting (anti-AASS antibody, Sigma-Aldrich, HPA020728) and enzyme activity assays.

*Protein expression and purification of the LOR domain*

The LOR domain from human AASS (NP_005754.2: residues 21-470) was cloned into a modified pET47 vector and expressed in *Escherichia coli* (BL21 LOBSTR cells). The cells were induced at OD 2.0 with 15 μM IPTG and cooled to 16°C overnight. The next day, the cells were lysed and protein was purified by IMAC and SEC to 95% purity. The protein was then concentrated to around 10 mg/ml and frozen with 10% glycerol.

*AASS/LOR forward activity assay in 96-well plate format*

LOR activity was measured in the forward direction using the substrates L-lysine, 2-oxoglutarate and NADPH. The products of the reaction are saccharopine and $NADP^+$. The established assay conditions for a 200μL reaction in a 96-well plate are 50 mM Hepes pH 7.4, 0.3 mM NADPH, 15 mM L-lysine and 0.25% Triton X-100. Recombinant LOR (0.1 μg) or Flp-In-293 AASS-Myc-DDK lysate (12 μg) were used as enzyme source and added as 50 μL/well, unless otherwise indicated. The plate was then preincubated for 10 minutes at 37°C. The reaction was started with the addition of 50 μL/well of 2-oxoglutarate to a final concentration of 1 mM or 10 mM as indicated. The stock solution containing 2-oxoglutarate was neutralized using KOH. The kinetics of the reaction were monitored as the decrease in absorption at  nm using a plate reader at 37°C. A substrate blank for correction of NADPH oxidation was included in all assays and subtracted. The millimolar extinction coefficient used for NADH/NADPH was 6220 $M^{-1}$ $cm^{-1}$. Steady-state kinetic data were analyzed by nonlinear regression analysis using GraphPad Prism 7 software and the Michaelis Menten or Hill equation was used. For conditions with substrate inhibition a modified Hill equation of LiCata and Allewell [1] for cooperative substrate binding as well as substrate inhibition was used, i.e. the velocity $v = (V_{max} + V_i([S]^x/K_i^x))/(1 + (K^{nH}/[S]^{nH}) + ([S]^x/K_i^x))$. The exponent x is a second Hill coefficient, which allows for the possibility that the substrate inhibition may also be cooperative [1], and by varying the value of x between 1 and 3, x = 2 gave the best fit for our values of the full-length enzyme.

*AASS/LOR reverse activity assay*
LOR activity was measured in the reverse direction using the substrates saccharopine and $NADP^+$. The assay conditions for a 200μL reaction in a 96-well plate are 100 mM Tris-HCl pH 9.0, 80 mM $NH_4Cl$, 1 mM $NADP^+$. Recombinant LOR at 0.0005 mg/mL (0.1 μg) or HEK-293 Flp-In AASS-Myc-DDK lysate (12 μg) were used as enzyme source and added as 50 μL/well, unless otherwise indicated. The plate was then preincubated for 10 minutes at 37°C. The reaction was started with the addition of 50 μL/well of saccharopine at a final concentration of 2 mM. The kinetics of the reaction were monitored as the increase in absorption at 340 nm using a plate reader at 37°C. A substrate blank for correction of $NADP^+$ reduction was included in all assays and subtracted.

*Virtual screening for LOR inhibitors*
The crystal structure of saccharopine dehydrogenase [$NAD^+$, L-lysine-forming] from *S. cerevisiae* with bound saccharopine and NADH (3UH1 [2]) was used to construct a homology model of the human LOR domain bound to saccharopine and NADPH [3]. Virtual screening was carried out using the Glide program of the Schrödinger Small-Molecule Drug Discovery Suite [4-6]. A library of more than 7 million commercially available drug-like compounds from the Molport collection was screened using the HTVS precision in Glide. In a second step, the top 100,000 hits from the HTVS screen were re-ranked using the standard precision (SP) in Glide. Finally, the top 10,000 compounds from the SP screen were re-ranked using the XP precision in Glide. Candidate hits were further filtered to eliminate compounds containing assay interfering chemical groups and promiscuous scaffolds [7], and were visually inspected in the context of the binding site. Based on structural and medicinal chemistry criteria, a final list of 127 compounds was selected for experimental validation using the LOR enzymatic assay and purchased through MolPort (Table S1; 69 from ChemBridge Corporation and 58 from ENAMINE Ltd.). Ultimately, 126 compounds were available, dissolved in DMSO and tested at 300 μM (1% DMSO final concentration), unless otherwise stated. $IC_{50}$ was determined for selected compounds using the GraphPad Prism 7 software.

*Repurchase of virtual screening compounds of interest*
After dose-response evaluation of virtual screening hits, compound **105** was purchased for further confirmation in AASS/LOR inhibition assays. Virtual screening ID MolPort-039-245-345 was obtained directly from ENAMINE Ltd. (#Z1457712064).

*SAR by catalog*
We followed up on the virtual screening hit compound 105 with SAR by catalog. We selected 37 analogs selected based on similarity and/or potency, and purchased this molecules through MolPort (33 from ENAMINE Ltd., 3 from UkrOrgSynthesis Ltd. Stock and 1 from Alinda Chemical, Ltd.).

*Synthesis of AASS inhibitors*

Methods for the re-synthesis of compound 105, its (*S*)- and (*R*)- isomers and a series of analogs is detailed below.

*GA1 cell line studies*

We used a previously generated CRISPR-Cas9-mediated *GCDH* KO HEK-293 cell line [8]. Cells were seeded in 6-well plates, and cultured for ~48h to reach 80-90% confluency. Cells then received fresh DMEM media with 10% FCS, Pen-Strep, 400μM L-carnitine and the compound at the indicated concentration (*S*-105 at 150 and 66.2μM). DMSO was used as vehicle control (0.45 and 0.2%, respectively). After 24h cells were harvested using trypsinization. Cell pellets were then sonicated in 200μL water after which 800μL acetonitrile was added to precipitate the protein. The supernatant was collected after centrifugation and internal labeled acylcarnitine standards were added (NSKB, Cambridge Isotope Laboratories). After drying under nitrogen, samples were butylated and analyzed by mass spectrometry as described in prior work [8, 9]. Protein pellets were dissolved in 25mM KOH and used for protein determination and normalization.

*Protein purification for crystallization*

Human AASS (amino acids 21-451; NP_005754) was cloned into a modified pET47 vector with an N-terminal His and Sumo-tag. The plasmid was transformed into LOBSTR-BL21(DE3). An overnight culture was used to inoculate 1 L of TB medium (1:1000) containing 50 μg/ml of kanamycin and grown at 37 °C until the cell density reached an OD600 of 1.0. The cells were then cooled to 16 °C and induced overnight with 15 μM isopropyl-β-D-thiogalactopyranoside (IPTG). The next day, cells were harvested and resuspended in tris-buffered saline (20 mM Tris pH 8.0, 250 mM NaCl) and lysed with a French pressure cell at 25K–30K p.s.i. After pelleting down the cellular debris, the lysate was loaded onto a column containing HisPurTM Ni-NTA Resin (Thermo Scientific) for immobilized metal affinity chromatography (IMAC) purification. The column was equilibrated with 7 column volumes of TBS supplemented with 40 mM imidazole pH 8.0 then washed with 7 column volumes of 20 mM Tris pH 8.0, 50 mM imidazole and adjusted to pH 7.5. The His tagged protein was eluted with 20 mM Tris pH 8.0 containing 150 mM NaCl, 150 mM imidazole and 10% glycerol. The proteolysis with Sumo protease to cleave off the tag was performed overnight at 4 °C in a 20 kDa cut-off dialysis cassette (Thermo Scientific) by dialyzing against no imidazole buffer (20 mM Tris, pH 8.0, 150 mM NaCl, 10% glycerol, 1 mM β-mercaptoethanol). Next day, the protein was incubated with Ni-NTA resin for an hour to bind the His-tagged protease and uncleaved protein, if any. The flow-through was collected and concentrated with a 30 kDa cut-off Amicon centrifugal filters (Millipore). This concentrated protein was purified further on Superdex 200 Increase gel filtration column (GE Lifescience) in a buffer of 20 mM Tris (pH 8.0), 150 mM NaCl, 10% glycerol.

For the longer human AASS construct, amino acids 21-470 were cloned into the same vector with an N-terminal His-Sumo tag. The protein was purified in the same manner as the shorter construct. However, the longer construct did not yield quality crystals, so we generated mutants that might be more favorable for crystallization. After analysis by the Surface Entropy Reduction server [10], we mutated five amino acids from two patches (residues 313-314 and 537-539 to alanine) that were predicted to hinder crystal packing. We then purified this construct in an identical manner.

*Crystallization and structure determination*

For both constructs, we screened crystallization conditions using commercial crystallization screens and identified conditions that gave initial crystals. We then optimized the crystals in larger hanging drops. For the short construct, the final protein crystallized in 0.2 M NaCl, 0.1 M HEPES, pH 7.5, 25% PEG 4000 (pro-complex D1). The longer construct crystallized in 11% PEG 3350, 0.1 M Magnesium Formate and 25 mM Calcium Acetate. Crystals were flash frozen in liquid nitrogen before collecting at the AMX (17-ID-1) beamline at NSLS II at Brookhaven National Laboratory. Data was processed using autoproc [11] and scaled with Aimless [12]. The structure was solved by molecular replacement, using the structure of the yeast saccharopine dehydrogenase (PDB 2QRJ) as a search model using the program Phaser [13] after making a homology model using Phenix Sculptor [14]. After molecular replacement, we performed multiples rounds of building using Phenix Autobuild [15]. The models were subsequently refined using Phenix [16] with rigid body refinement and multiple rounds of simulated annealing, minimization, atomic displacement parameter (ADP or B-factor) refinement and TLS refinement (determined using the TLSMD server) [17, 18] with interspersed manual adjustments using Coot [19]. All structural figures were made with Pymol [20]. For the longer construct, we solved the structure by molecular replacement using our short construct structure as a search model. The model was built similarly, by iterative rounds of manual building in Coot and in Phenix Autobuild to complete the missing regions such as the C-terminal extension.

*Animal experiments*

The plasma lysine data were re-analyzed from a previous work . [9] For the high lysine exposure, we generated *Gcdh* KO, *Aass* KO, *Gcdh*/*Aass* DKO as well as WT controls on a mixed, but well-defined background (F2 of C57BL/6J x 129S2/SvPasCrl). The first cohorts of animals (21 mice) were exposed to high lysine through diet (TD.04412, L-lysine added to a standard rodent diet to a total amount of 4.7% w/w) and drinking water (4.7% L-lysine w/v) at 4 weeks of age [21, 22]. Since we noted an aversion of the mice to the high lysine diet, we also exposed one cohort of 10 mice to high lysine by increasing the dose in drinking water (9.4% L-lysine w/v) and feeding regular chow diet. The latter protocol worked better as demonstrated by normal food and water intake in all animals. Importantly, we were able to reach the target daily dose of lysine (329 mg [21, 22]). All mice in the study were evaluated daily for body weight loss and motor deficits using a simple quantitative neurological scale developed in rats [23, 24] and used previously to monitor disease progression in the GA1 mouse model [25]. This evaluation assesses degree of hindlimb dystonia, gait abnormalities, recumbency and paralysis, and forepaw grasping [23, 24]. After a maximum of 3 days of high lysine exposure, the mice were euthanized and organs and plasma collected for histopathology and clinical biochemistry, respectively. Mice were euthanized at any earlier time-point if neurological disease progressed faster and/or body weight loss was >20% of the pre-study weight. Plasma amino acids were measured as described . [26]

### Synthesis of AASS inhibitors

Scheme 1. Synthesis of (*S*)-, (*R*)- and racemic compound #105[a]

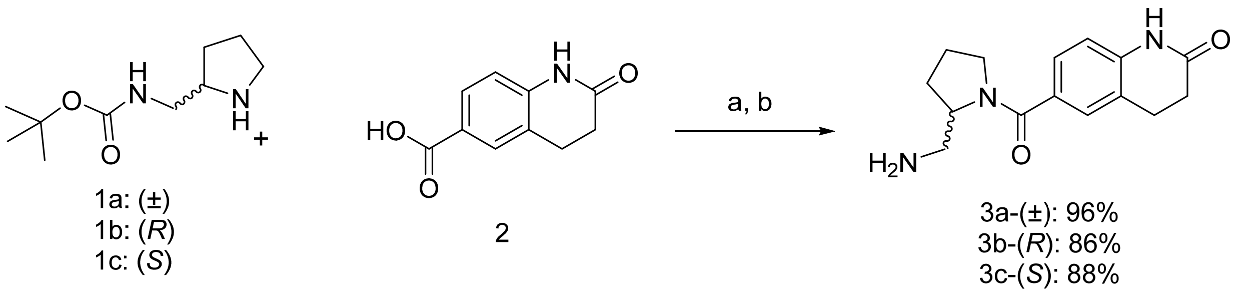

[a]Reagents and conditions: (a) EDC, HOBt, TEA, DMF, 25°C, 16 h (b) 4M HCl, dioxane, DCM, rt, 12 h, yield: 96-86%

Scheme 2. Synthesis of compound #165-169 and 172[a]

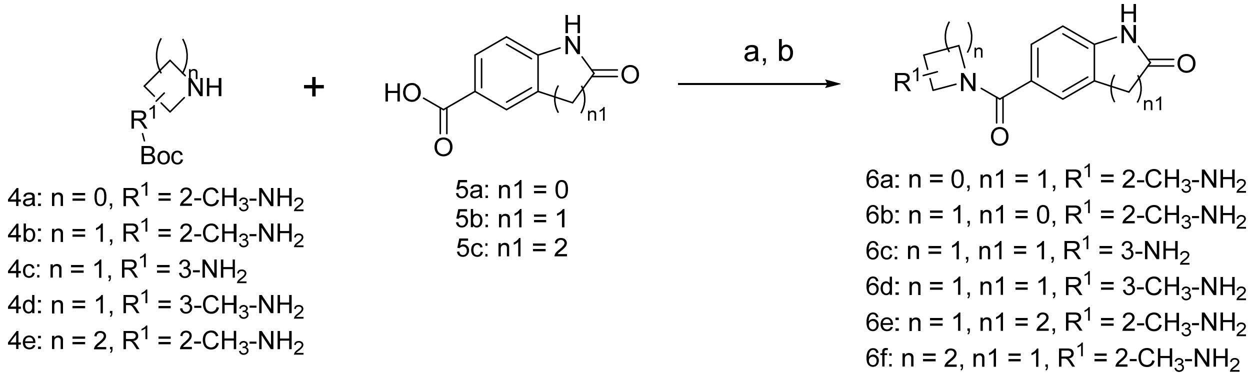

Compound #165 = 6f, compound #166 = 6c, compound #167 = 6a, compound #168 = 6d, compound #169 = 6b and compound #172 = 6e.

[a]Reagents and conditions: (a) HATU, $Et_3N$, DMF, rt, 12 h (b) 4M HCl, dioxane, DCM, rt, 12 h, yield: 58-17%

Scheme 3. Synthesis of compound #170 and 171[a]

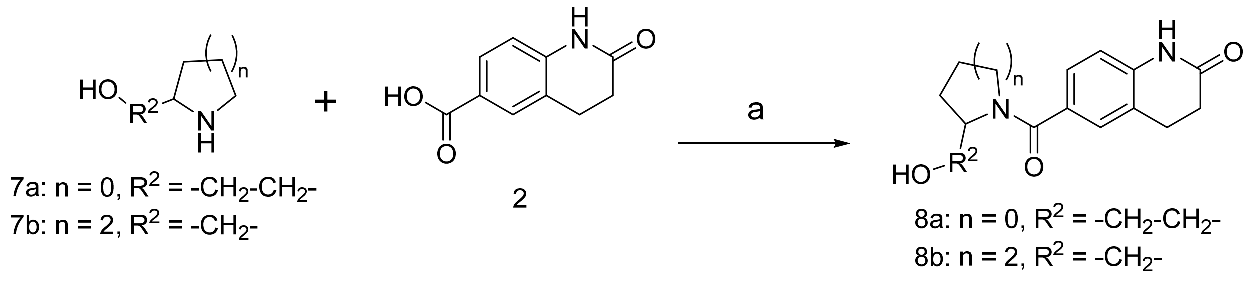

Compound #170 = 8a and compound #171 = 8b.

[a]Reagents and conditions: (a) HATU, $Et_3N$, DMF, rt, 12 h, yield: 20-17%

Scheme 4. Synthesis of compound #173 and 174[a]

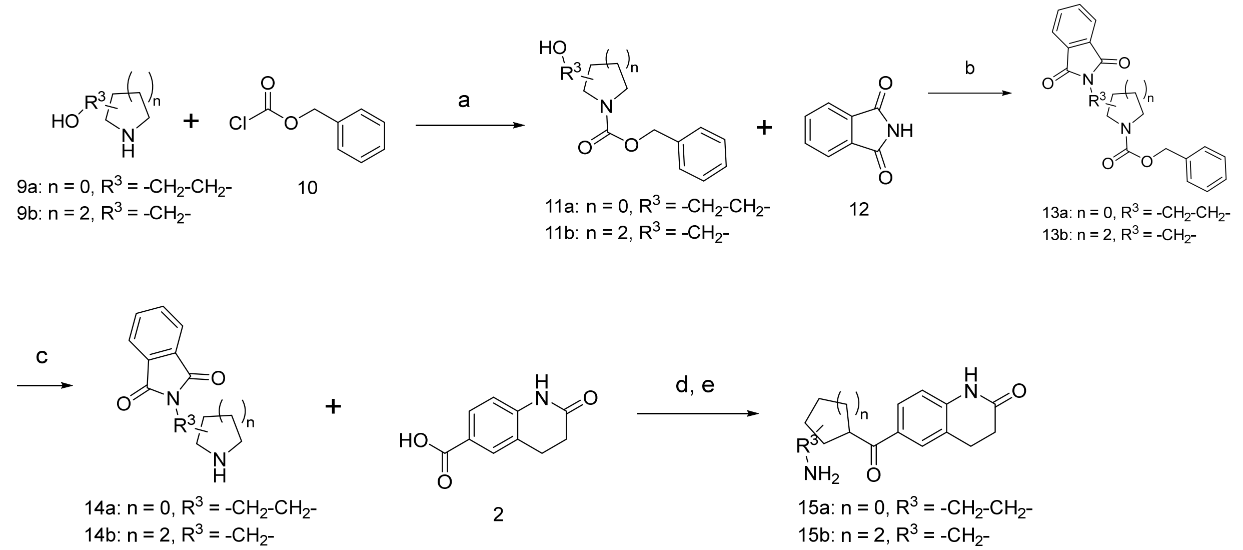

Compound #173 = 15b and compound #174 = 15a.

[a]Reagents and conditions: (a) $K_2CO_3$, THF/$H_2O$, rt, 16 h (b) DEAD, $PPh_3$, THF, 0$^o$C to rt, 16 h (c) Pd/C, $H_2$, MeOH, rt, 16 h (d) HATU, $Et_3N$, DMF, rt, 12 h (e) hydrazine hydrate, 50-60%, EtOH, rt, 16 h, yield: 50-10%

### Materials

Materials and Methods. $^1$H spectra were acquired on a Bruker DRX-600 spectrometer at 600 MHz for $^1$H. Thin layer chromatography (TLC) was performed on silica coated aluminum sheets (thickness 200 μm) or alumina coated (thickness 200 µm) aluminum sheets supplied by Sorbent Technologies, and column chromatography was carried out on Teledyne ISCO combiflash equipped with a variable wavelength detector and a fraction collector using a RediSep Rf high performance silica flash columns by Teledyne ISCO. LCMS/HPLC analysis for purity determination and HRMS was conducted on an Agilent Technologies G1969A high-resolution API-TOF mass spectrometer attached to an Agilent Technologies 1200 HPLC system. Samples were ionized by electrospray ionization (ESI) in positive mode. Chromatography was performed on a 2.1 × 150 mm Zorbax 300SBC18 5-µm column with water containing 0.1% formic acid as solvent A and acetonitrile containing 0.1% formic acid as solvent B at a flow rate of 0.4 mL/min. The gradient program was as follows: 1% B (0−1 min), 1−99% B (1−4 min), and 99% B (4−8 min). The temperature of the column was held at 50°C for the entire analysis. The purity of all the compounds was ≥95%. The chemicals and reagents were purchased from Combi-Blocks, Enamine, Oakwood Chemical, Sigma-Aldrich and Acros Organics. All solvents were purchased in anhydrous from Acros Organics and used without further purification.

#### Experimental Procedure

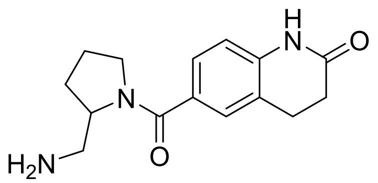

**6-(2-(aminomethyl)pyrrolidine-1-carbonyl)-3,4-dihydroquinolin-2(1H)-one hydrochloride (Racemic compound #105, 3a)**: General procedure A: To a solution of 2-oxo-3,4-dihydro-1H-quinoline-6-carboxylic acid (97.2 mg, 0.51 mmol), tert-butyl (pyrrolidin-2-ylmethyl)carbamate (112.0 mg, 0.56 mmol) and HOBt (85.6 mg, 0.56 mmol) in DMF (5 mL) was added, EDC (107.0 mg, 0.56 mmol) and triethylamine (0.08 mL, 0.56 mmol) at 0ºC under argon.  The reaction was stirred at 0 ºC to rt for 16 hours. Water (10 mL) was added and the aqueous layer was extracted with ethyl acetate (3 × 15 mL). The combined organic extracts were washed with brine, dried over anhydrous $Na_2SO_4$, filtered, and concentrated. The residue was purified by column chromatography on silica gel (100% dichloromethane to 5% methanol in dichloromethane) to obtain a yellow solid intermediate, tert-butyl

((1-(2-oxo-1,2,3,4-tetrahydroquinoline-6-carbonyl)pyrrolidin-2-yl)methyl)carbamate: LCMS (ESI): m/z $[M + H]^+$ = 374.2105. The BOC intermediate was stirred with HCl, 4M in dioxane (0.4 mL, 2.0 mmol) and dichloromethane (2 mL) at 25 °C for 12 hours. After 12 hours, the solvent was removed in vacuo. Toluene (3 mL) was added and removed *in vacuo.* The crude product was washed with dichloromethane and diethyl ether to afford 6-(2-(aminomethyl)pyrrolidine-1-carbonyl)-3,4-dihydroquinolin-2(1*H*)-one hydrochloride (yield: 14.8 mg; 96%) as a white solid. $^1$H NMR (DMSO-$d_6$, 400 MHz) δ 10.27 (s, 1H), 7.87 (broad s, 2H), 7.39–7.36 (m, 2H), 6.87 (d, *J* = 8.1 Hz, 1H), 5.75 (s, 1H), 4.34–4.25 (m, 1H), 3.57–3.51 (m, 1H), 3.43–3.37 (m, 1H), 3.14–2.88 (m, 4H), 2.47 (t, *J* = 7.6 Hz, 2H), 2.07–2.03 (m, 1H), 1.87–1.73 (m, 2H); HRMS (ESI): m/z $[M + H]^+$ calculated for $C_{15}H_{20}N_3O_2$ 274.1550; found 274.1479; purity >95%.

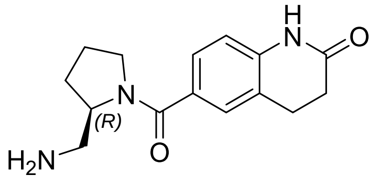

**(*R*)-6-(2-(aminomethyl)pyrrolidine-1-carbonyl)-3,4-dihydroquinolin-2(1H)-one hydrochloride (Compound #*R*-105, 3b)**: The compound synthesis was according to the procedure A. The intermediate, tert-butyl (*R*)-((1-(2-oxo-1,2,3,4-tetrahydroquinoline-6-carbonyl)pyrrolidin-2-yl)methyl)carbamate: LCMS (ESI): m/z $[M + H]^+$ = 374.2067. The key product, (*R*)-6-(2-(aminomethyl)pyrrolidine-1-carbonyl)-3,4-dihydroquinolin-2(1*H*)-one hydrochloride (yield: 62.8 mg; 88%). $^1$H NMR (MeOD-$d_4$, 400 MHz) 7.47 (broad s, 1H), 7.44 (broad s, 1H), 6.96–6.95 (d, *J* = 7.6 Hz, 1H), 4.45 (broad s, 1H), 3.66 (broad s, 1H), 3.60 (broad s, 1H), 3.24 (broad s, 2H), 3.02 (broad s, 2H), 2.64-2.60 (t, *J* = 6.4, 7.2 Hz, 2H), 2.30 (broad s, 1H), 1.99 (broad s, 1H), 1.84 (broad s, 2H); HRMS (ESI): m/z $[M + H]^+$ calculated for $C_{15}H_{20}N_3O_2$ 274.1550; found 274.1571; purity >95%.

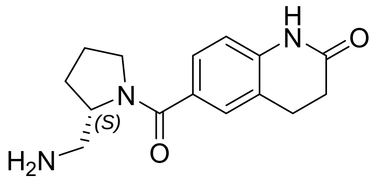

**(*S*)-6-(2-(aminomethyl)pyrrolidine-1-carbonyl)-3,4-dihydroquinolin-2(1H)-one hydrochloride (Compound #*S*-105, 3c)**: The compound synthesis was according to the procedure A. The intermediate, tert-butyl (*S*)-((1-(2-oxo-1,2,3,4-tetrahydroquinoline-6-carbonyl)pyrrolidin-2-yl)methyl)carbamate: LCMS (ESI): m/z $[M + H]^+$ = 374.2251. The key product, (*S*)-6-(2-(aminomethyl)pyrrolidine-1-carbonyl)-3,4-dihydroquinolin-2(1H)-one hydrochloride (yield: 86.6 mg; 86%). $^1H$ NMR (MeOD-$d_4$, 400 MHz) 7.47 (broad s, 1H), 7.44 (broad s, 1H), 6.96–6.95 (d, $J$ = 7.2 Hz 1H), 4.45 (broad s, 1H), 3.66 (broad s, 1H), 3.60 (broad s, 1H), 3.24 (broad s, 2H), 3.02 (broad s, 2H), 2.62 (broad s, 2H), 2.30 (broad s, 1H), 1.99 (broad s, 1H), 1.84 (broad s, 2H); HRMS (ESI): m/z $[M + H]^+$ calculated for $C_{15}H_{20}N_3O_2$ 274.1550; found 274.1591; purity >95%.

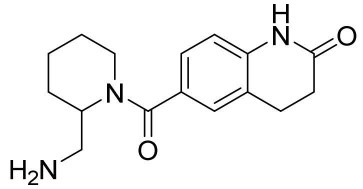

**6-(2-(aminomethyl)piperidine-1-carbonyl)-3,4-dihydroquinolin-2(1H)-one (Compound #165, 6f)**: General procedure B: A solution of tert-butyl (piperidin-2-ylmethyl)carbamate (0.112 g, 1 eq, 0.523 mmol), 2-oxo-1,2,3,4-tetrahydroquinoline-6-carboxylic acid (0.1 g, 1 eq, 0.523 mmol), HATU (0.298 g, 1.5 eq, 0.785 mmol) and triethylamine (0.219 mL, 3.0 eq, 1.57 mmol) in DMF (5mL) was stirred at 25 °C for 16 hour. After 16 hours, the reaction was extracted with ethyl acetate and water. The ethyl acetate layer was collected, dried with sodium sulfate and purified by the normal phase (100% hexane to 100% ethyl acetate) to obtain tert-butyl ((1-(2-oxo-1,2,3,4-tetrahydroquinoline-6-carbonyl)piperidin-2-yl)methyl)carbamate. The impure Boc intermediate fractions were collected and concentrated to obtain an oil residue: LCMS (ESI): m/z $[M + H]^+$ = 389.3796. The residue was stirred with HCl, 4M in dioxane (0.39 mL, 1.57 mmol) and $CH_2Cl_2$ (2 mL) at 25 °C for 12 hours. After 16 hours, the reaction was neutralized with 7N ammonia in methanol and purified by the normal phase (100% dichloromethane to 20% methanol/0.1% ammonium hydroxide in dichloromethane) or the reversed phase (5% methanol in 0.1% trifluoroacetic acid/water to 40% methanol in 0.1% trifluoroacetic acid/water) to afford 6-(2-(aminomethyl)piperidine-1-carbonyl)-3,4-dihydroquinolin-2(1H)-one (yield: 0.0772 g; 52%). $^1H$ NMR (MeOD-$d_4$, 600 MHz) δ 7.37 (broad s, 1H), 7.34 (d, $J$ = 8.03 Hz, 1H), 6.97 (d, $J$ = 8.10 Hz, 1H), 4.97 (broad s, 1H), 3.77 (broad s, 1H), 3.64-3.53 (m, 1H), 3.21 (broad s, 1H), 3.17 (dd, $J$ = 13.31, 4.59 Hz, 1H), 3.02 (t, $J$ = 7.56 Hz, 2H),

2.62 (t, *J* = 15.26 Hz, 2H), 2.00-1.56 (m, 6H). HRMS (ESI): m/z $[M + H]^+$ calcd for $C_{16}H_{21}N_3O_2^+$: 288.1710, found: 288.1916; purity >95%.

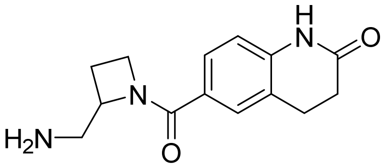

**6-(2-(aminomethyl)azetidine-1-carbonyl)-3,4-dihydroquinolin-2(1H)-one (Compound #167, 6a)**: The compound synthesis was according to the procedure B. The intermediate, tert-butyl ((1-(2-oxo-1,2,3,4-tetrahydroquinoline-6-carbonyl)azetidin-2-yl)methyl)carbamate: LCMS (ESI): m/z $[M + H]^+$ = 360.3128. The key product, 6-(2-(aminomethyl)azetidine-1-carbonyl)-3,4-dihydroquinolin-2(1H)-one (yield: 0.0791 g; 58%). $^1$H NMR (MeOD-$d_4$, 600 MHz) δ 7.73 (m, 1H), 7.56-7.53 (m, 1H), 6.95-6.93 (m, 1H), 4.74 (d, *J* = 118.68 Hz, 1H), 4.37 (d, *J* = 74.70 Hz, 1H), 3.62-3.56 (m, 1H), 3.44-3.40 (m, 1H), 3.01 (t, *J* = 6.21 Hz, 2H), 2.61 (t, *J* = 7.47 Hz, 2H), 2.58-2.39 (m, 1H), 2.27-2.19 (m, 1H). HRMS (ESI): m/z $[M + H]^+$ calcd for $C_{14}H_{17}N_3O_2^+$: 260.1394, found: 260.1420; purity >95%.

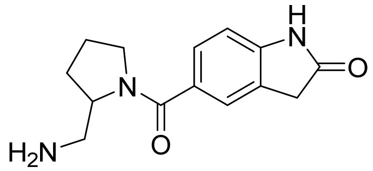

**5-(2-(aminomethyl)pyrrolidine-1-carbonyl)indolin-2-one (Compound #169, 6b)**: The compound synthesis was according to the procedure B. The intermediate, tert-butyl ((1-(2-oxoindoline-5-carbonyl)pyrrolidin-2-yl)methyl)carbamate: LCMS (ESI): m/z $[M + H]^+$ = 360.3795. The key product, 5-(2-(aminomethyl)pyrrolidine-1-carbonyl)indolin-2-one (yield: 0.0587 g; 40%). $^1$H NMR (MeOD-$d_4$, 600 MHz) δ 7.52 (d, *J* = 8.07 Hz, 2H), 6.99 (dd, *J* = 4.09 Hz, 1H), 4.47-4.45 (m, 1H), 3.72-3.67 (m, 1H), 3.60 (broad s, 3H), 3.30-3.27 (m, 1H), 3.24-3.22 (m, 1H), 2.32-2.29 (m, 1H), 2.02-1.96 (m, 1H), 1.89-

1.81 (m, 1H). HRMS (ESI): m/z $[M + H]^+$ calcd for $C_{14}H_{17}N_3O_2^+$: 260.1394, found: 260.1431; purity >95%.

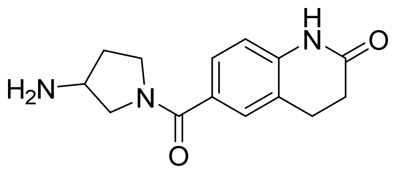

**6-(3-aminopyrrolidine-1-carbonyl)-3,4-dihydroquinolin-2(1H)-one (Compound #166, 6c)**: The compound synthesis was according to the procedure B. The intermediate, tert-butyl (1-(2-oxo-1,2,3,4-tetrahydroquinoline-6-carbonyl)pyrrolidin-3-yl)carbamate: LCMS (ESI): m/z $[M + H]^+$ = 360.3628. The key product, 6-(3-aminopyrrolidine-1-carbonyl)-3,4-dihydroquinolin-2(1H)-one (yield: 0.0676 g; 50%). $^1$H NMR (MeOD-$d_4$, 600 MHz) δ 7.44 (broad s, 2H), 6.96 (d, $J$ = 8.11 Hz, 1H), 4.03 (broad s, 1H), 3.96 (dd, $J$ = 6.20 Hz, 1H), 3.83-3.59 (m, 3H), 3.01 (t, $J$ = 7.51 Hz, 2H), 2.61 (t, $J$ = 15.07 Hz, 2H), 2.40 (broad s, 1H), 2.13 (broad s, 1H). HRMS (ESI): m/z $[M + H]^+$ calcd for $C_{14}H_{17}N_3O_2^+$: 260.1394, found: 260.1770; purity >95%.

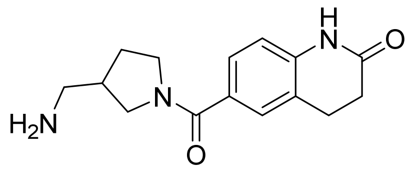

**6-(3-(aminomethyl)pyrrolidine-1-carbonyl)-3,4-dihydroquinolin-2(1H)-one (Compound #168, 6d)**: The compound synthesis was according to the procedure B. The intermediate, tert-butyl ((1-(2-oxo-1,2,3,4-tetrahydroquinoline-6-carbonyl)pyrrolidin-3-yl)methyl)carbamate: LCMS (ESI): m/z $[M + H]^+$ = 374.2082. The key product, 6-(3-(aminomethyl)pyrrolidine-1-carbonyl)-3,4-dihydroquinolin-2(1H)-one (yield: 0.0384 g; 27%). $^1$H NMR (MeOD-$d_4$, 400 MHz) δ 7.43 (broad s, 2H), 6.96 (broad s, 1H), 3.88 (broad s, 1H), 3.77 (broad s, 1H), 3.69 (broad s, 1H), 3.39 (broad s, 1H), 3.13 (broad s, 1H), 3.02 (broad s, 3H), 2.64 (broad s, 3H), 2.23 (broad s, 1H), 1.81 (broad s, 1H). HRMS (ESI): m/z $[M + H]^+$ calcd for $C_{15}H_{19}N_3O_2^+$: 274.1550, found: 274.1558; purity >95%.

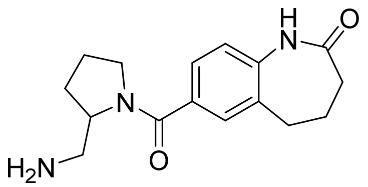

**7-(2-(aminomethyl)pyrrolidine-1-carbonyl)-1,3,4,5-tetrahydro-2H-benzo[b]azepin-2-one (Compound #172, 6e)**: The compound synthesis was according to the procedure B. The intermediate, tert-butyl ((1-(2-oxo-2,3,4,5-tetrahydro-1H-benzo[b]azepine-7-carbonyl)pyrrolidin-2-yl)methyl)carbamate: LCMS (ESI): m/z $[M + H]^+$ = 388.0799. The key product, 7-(2-(aminomethyl)pyrrolidine-1-carbonyl)-1,3,4,5-tetrahydro-2H-benzo[b]azepin-2-one (yield: 0.0308 g; 22%). $^1$H NMR (MeOD-$d_4$, 600 MHz) δ 7.52 (s, 1H), 7.50 (d, $J$ = 8.07 Hz, 1H), 7.11 (d, $J$ = 8.02 Hz, 1H), 4.48-4.43 (m, 1H), 3.67-3.63 (m, 1H), 3.60-3.56 (m, 1H), 3.29-3.25 (m, 1H), 3.22-3.20 (m, 1H), 2.85 (t, $J$ = 7.08, 7.25 Hz, 2H), 2.35-2.26 (m, 5H), 2.02-1.97 (m, 1H), 1.90-1.80 (m, 2H). HRMS (ESI): m/z $[M + H]^+$ calcd for $C_{16}H_{21}N_3O_2^+$: 288.1707, found: 288.2074; purity >95%.

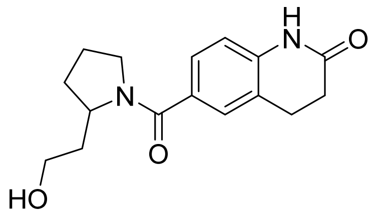

**6-(2-(2-hydroxyethyl)pyrrolidine-1-carbonyl)-3,4-dihydroquinolin-2(1H)-one (Compound #170, 8a)**: General procedure C: A solution of 2-(pyrrolidin-2-yl)ethan-1-ol (0.1 g, 1.0 eq, 0.661 mmol), 2-oxo-1,2,3,4-tetrahydroquinoline-6-carboxylic acid (0.126 g, 1.0 Eq, 0.661 mmol), HATU (0.377 g, 1.5 eq, 0.992 mmol) and triethylamine (0.277 mL, 3.0 eq, 1.98 mmol) in DMF (5 mL) was stirred at 25 °C for 16 hour. After 16 hours, the reaction was purified by the reversed phase (5% methanol in 0.1%

trifluoroacetic acid/water to 40% methanol in 0.1% trifluoroacetic acid/water) to afford 6-(2-(2-hydroxyethyl)pyrrolidine-1-carbonyl)-3,4-dihydroquinolin-2(1H)-one (yield: 0.0389 g; 20%). $^{1}$H NMR (MeOD-*d*$_4$, 600 MHz) δ 7.88 (broad s, 1H), 7.87 (broad s, 1H), 6.94 (d, *J* = 8.08 Hz, 1H), 4.49-4.45 (m, 1H), 4.43-4.39 (m, 1H), 3.74-3.69 (m, 1H), 3.40-3.31 (m, 2H), 3.01 (t, *J* = 7.49, 7.68 Hz, 2H), 2.61 (t, *J* = 7.31, 7.90 Hz, 2H), 2.37-3.31 (m, 1H), 2.29-2.18 (m, 2H), 2.16-2.02 (m, 2H), 1.81-1.74 (m, 1H). HRMS (ESI): m/z [M + H]$^+$ calcd for $C_{16}H_{20}N_2O_3^+$: 289.1547, found: 289.1783; purity >95%.

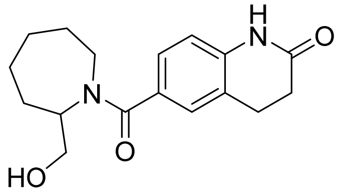

**6-(2-(hydroxymethyl)azepane-1-carbonyl)-3,4-dihydroquinolin-2(1H)-one (Compound #171, 8b)**: The compound synthesis was according to the procedure C (yield: 0.0406 g; 17%). $^{1}$H NMR (MeOD-*d*$_4$, 400 MHz) δ 7.96 (d, *J* = 9.62 Hz, 1H), 7.26 (broad s, 1H), 6.97 (d, *J* = 8.59 Hz, 1H), 4.55-4.53 (m, 1H), 4.45-4.40 (m, 1H), 3.79 (broad s, 1H), 3.42-3.30 (m, 2H), 3.11-2.96 (m, 2H), 2.69-2.61 (m, 2H), 2.11-1.72 (m, 7H), 1.36 (m, 1H). HRMS (ESI): m/z [M + H]$^+$ calcd for $C_{17}H_{22}N_2O_3^+$: 303.1703, found: 303.2439; purity >95%.

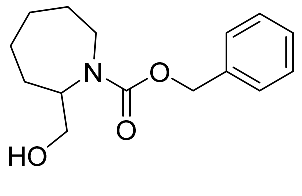

**Benzyl 2-(hydroxymethyl)azepane-1-carboxylate (11b)**: General procedure D: A solution of azepan-2-ylmethanol (0.1 g, 1 eq, 0.774 mmol) and potassium carbonate (0.481 g, 4.5 eq, 3.48 mmol) in THF (3 mL)/Water (1 mL) was vividly stirred at 25 °C. Liquid of benzyl chloroformate (0.132 g, 0.109 mL, 1.0 Eq, 0.774 mmol) was dropwised in the reaction. The reaction was then stirred at 25 °C for 16 hours. After 16 hours, the reaction mixture was extracted with ethyl acetate and water. The ethyl acetate layer was collected, dried with sodium sulfate and purified by the normal phase (100% hexane to 50% ethyl acetate in hexane) to afford benzyl 2-(hydroxymethyl)azepane-1-carboxylate (yield: 0.1853 g; 91%). $^{1}$H NMR (Chloroform-*d*, 600 MHz) δ 7.37 (broad s, 4H), 7.33-7.32 (m, 1H), 5.20-5.14 (m, 2H), 4.23-4.10 (m, 1H), 3.85 (dd, *J* = 14.25, 14.72, 39.46 Hz, 1H), 3.64 (dd, *J* = 3.72, 3.76, 7.26 Hz, 1H), 3.57-3.51 (m, 1H), 2.88-2.83 (m, 1H), 2.68 (broad s, 1H), 2.06-1.99 (m, 1H), 1.85-1.81 (m, 1H), 1.73 (d, *J* = 14.26 Hz, 1H), 1.60-1.47 (m, 1H), 1.40-1.21 (m, 3H). LCMS (ESI): m/z $[M + H]^+$ = found: 264.2615.

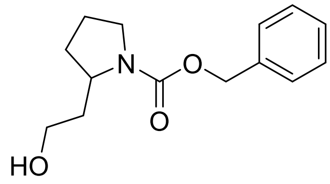

**Benzyl 2-(2-hydroxyethyl)pyrrolidine-1-carboxylate (11a)**: The compound synthesis was according to the procedure D (yield: 0.1942 g; 87%). $^{1}$H NMR (Chloroform-*d*, 600 MHz) δ 7.38-7.35 (m, 4H), 7.33-7.32 (m, 1H), 4.21 (broad s, 1H), 3.66-3.58 (m, 4H), 3.45-3.39 (m, 2H), 2.05-1.97 (m, 1H), 1.91 (broad s, 2H), 1.71 (broad s, 1H), 1.67 (broad s, 1H), 1.56 (broad s, 1H). LCMS (ESI): m/z $[M + H]^+$ = 250.2664.

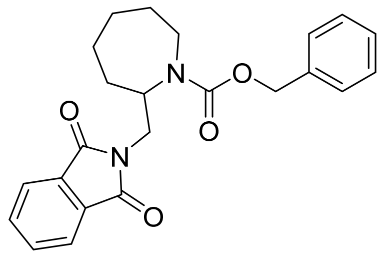

**Benzyl 2-((1,3-dioxoisoindolin-2-yl)methyl)azepane-1-carboxylate (13b):** General procedure E: A solution of benzyl 2-(hydroxymethyl)azepane-1-carboxylate (0.204 g, 1 eq, 0.774 mmol), isoindoline-1,3-dione (0.182 g, 1.6 eq, 1.24 mmol) and triphenylphosphine (0.325 g, 1.6 eq, 1.24 mmol) in anhydrous THF (10 mL) at 25 °C for 1 hour. Liquid of diethyl azodicarboxylate (DEAD), 40% solution in toluene (0.55 mL, 1.6 eq, 1.24 mmol) was dropwised in the reaction at 0 °C. The reaction was then warmed up to 25 °C and stirred for 16 hours. After 16 hours, the reaction was quenched with water and extracted with ethyl acetate. The ethyl acetate layer was collected, dried with sodium sulfate and purified by the normal phase (100% hexane to 50% ethyl acetate in hexane) to afford benzyl 2-((1,3-dioxoisoindolin-2-yl)methyl)azepane-1-carboxylate (yield: 0.1612 g; 53%). $^{1}$H NMR (Chloroform-*d*, 600 MHz) δ 7.74-7.72 (m, 1H), 7.66-7.64 (m, 1H), 7.63-7.60 (m, 2H), 7.24-7.20 (m, 3H), 7.13 (d, *J* = 7.55 Hz, 1H), 7.09 (t, *J* = 3.49, 3.70 Hz, 1H), 4.70 (dd, *J* = 12.46, 12.47, 248.28 Hz, 1H), 4.84 (quartet, *J* = 7.19, 12.66 Hz, 1H), 4.54-4.39 (m, 1H), 3.08-3.52 (m, 3H), 2.88-2.82 (m, 1H), 2.17-2.08 (m, 1H), 1.80-1.76 (m, 2H), 1.67 (t, *J* = 12.58, 13.05 Hz, 1H)), 1.56-1.41 (m, 1H), 1.33-1.16 (m, 3H). LCMS (ESI): m/z $[M + H]^+$ = 393.4289

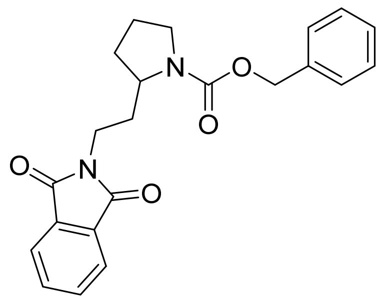

**Benzyl 2-(2-(1,3-dioxoisoindolin-2-yl)ethyl)pyrrolidine-1-carboxylate (13a)**: The compound synthesis was according to the procedure E (yield: 0.1558; 46%). $^{1}$H NMR (Chloroform-*d*, 600 MHz) δ 7.75 (broad s, 2H), 7.71 (m, 2H), 7.24 (broad s 3H), 7.11 (broad s 2H), 4.91 (m, 2H), 3.78 (broad s, 1H), 3.62 (m, 1H), 3.57 (m, 1H), 3.32 (m, 2H), 2.00 (m, 2H), 1.87 (m, 1H), 1.79 (m, 2H), 1.67-1.60 (m, 1H). LCMS (ESI): m/z $[M + H]^+$ = found: 379.5315.

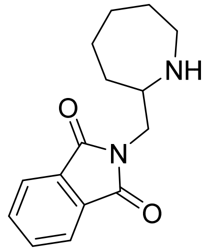

**2-(azepan-2-ylmethyl)isoindoline-1,3-dione (14b)**: General procedure F: A solution of benzyl 2-((1,3-dioxoisoindolin-2-yl)methyl)azepane-1-carboxylate (0.1612 g, 1 eq), Pd/C (0.05 g) and hydrogen gas in methanol (5 mL) was stirred at 25 °C for 16 hour. After 16 hours, the reaction was filtered with celite pad and the filtrate was monitored by LCMS. LCMS (ESI): m/z $[M + H]^+$ = 259.3132. The filtrate was concentrated to obtain an oil residue used for the next step without chromatographic purification.

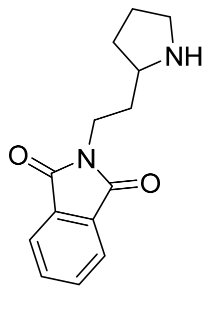

2-(2-(pyrrolidin-2-yl)ethyl)isoindoline-1,3-dione **(14a)**: The compound synthesis was according to the procedure F. LCMS (ESI): m/z $[M + H]^+$ = 245.2682.

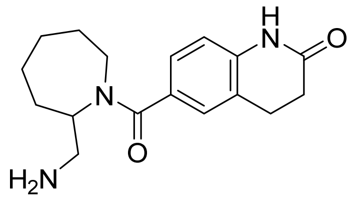

**6-(2-(aminomethyl)azepane-1-carbonyl)-3,4-dihydroquinolin-2(1H)-one (Compound #173, 15b)**: General procedure G: A solution of 2-(azepan-2-ylmethyl)isoindoline-1,3-dione (0.1061 g, 1 eq, 0.4107 mmol), 2-oxo-1,2,3,4-tetrahydroquinoline-6-carboxylic acid (0.07852 g, 1.0 eq, 0.4107 mmol), HATU (0.2342 g, 1.5 eq, 0.6161 mmol) and triethylamine (0.18 mL, 3.0 eq, 1.232 mmol) in DMF (5 mL) was stirred at 25 °C for 16 hour. The reaction was extracted with ethyl acetate and water. The ethyl acetate layer was collected, dried with sodium sulfate and purified by the normal phase (100% hexane to 100% ethyl acetate) to afford the impure intermediate, 2-((1-(2-oxo-1,2,3,4-tetrahydroquinoline-6-carbonyl)azepan-2-yl)methyl)isoindoline-1,3-dione. LCMS (ESI): m/z $[M + H]^+$ = 432.4221. A solution of 2-((1-(2-oxo-1,2,3,4-tetrahydroquinoline-6-carbonyl)azepan-2-yl)methyl)isoindoline-1,3-dione (0.3556 g, 1 eq, 0.8242 mmol) and hydrazine hydrate, 50-60% (0.1 mL) in ethanol (5 mL) was stirred at 25 °C for 16 hour. After 16 hours, the reaction was neutralized with HCl 4M in dioxane to be pH 7.0, and the reaction mixture was then purified by the reversed phase (5% methanol in water to 40% methanol in water) to afford 6-(2-(aminomethyl)azepane-1-carbonyl)-3,4-dihydroquinolin-2(1H)-one (yield: 0.1042 g; 50%). $^1$H NMR (MeOD-$d_4$, 400 MHz) δ 7.76 (broad s, 1H), 7.35 (d, $J$ = 11.41 Hz, 1H), 6.95 (broad s, 1H), 4.78 (broad s, 1H), 3.62-3.57 (m, 2H), 3.24 (broad s, 1H), 3.11-2.95 (m, 3H), 2.62 (broad s, 2H), 2.18-1.40(m, 8H). HRMS (ESI): m/z $[M + H]^+$ calcd for $C_{17}H_{23}N_3O_2^+$: 302.1863, found: 302.2392; purity >95%.

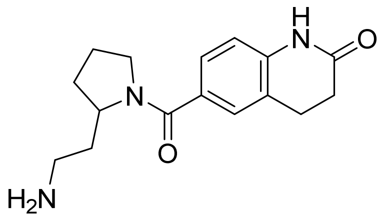

**6-(2-(2-aminoethyl)pyrrolidine-1-carbonyl)-3,4-dihydroquinolin-2(1H)-one (Compound #174, 15a):** The compound synthesis was according to the procedure G. The intermediate, 2-(2-(1-(2-oxo-1,2,3,4-tetrahydroquinoline-6-carbonyl)pyrrolidin-2-yl)ethyl)isoindoline-1,3-dione: LCMS (ESI): m/z $[M + H]^+$ = 418.3244. The key product, 6-(2-(2-aminoethyl)pyrrolidine-1-carbonyl)-3,4-dihydroquinolin-2(1H)-one (yield: 0.0105 g; 10%). $^1$H NMR (MeOD-$d_4$, 600 MHz) δ 7.40 (broad s, 1H), 7.37 (d, $J$ = 8.05 Hz, 1H), 6.92 (d, $J$ = 8.02 Hz, 1H), 4.34 (m, 1H), 3.64-3.60 (m, 1H), 3.58-3.43 (m, 1H), 3.05 (m, 1H), 2.99 (m, 3H), 2.59 (t, $J$ = 7.31, 7.45 Hz, 2H), 2.22-2.18 (m, 1H), 2.15-2.11 (m, 1H), 2.04-1.97 (m, 1H), 1.92-1.86 (m, 1H), 1.84-1.80 (m, 1H), 1.75-1.67 (m, 1H). HRMS (ESI): m/z $[M + H]^+$ calcd for $C_{16}H_{21}N_3O_2^+$: 288.1707, found: 288.2109; purity >95%.

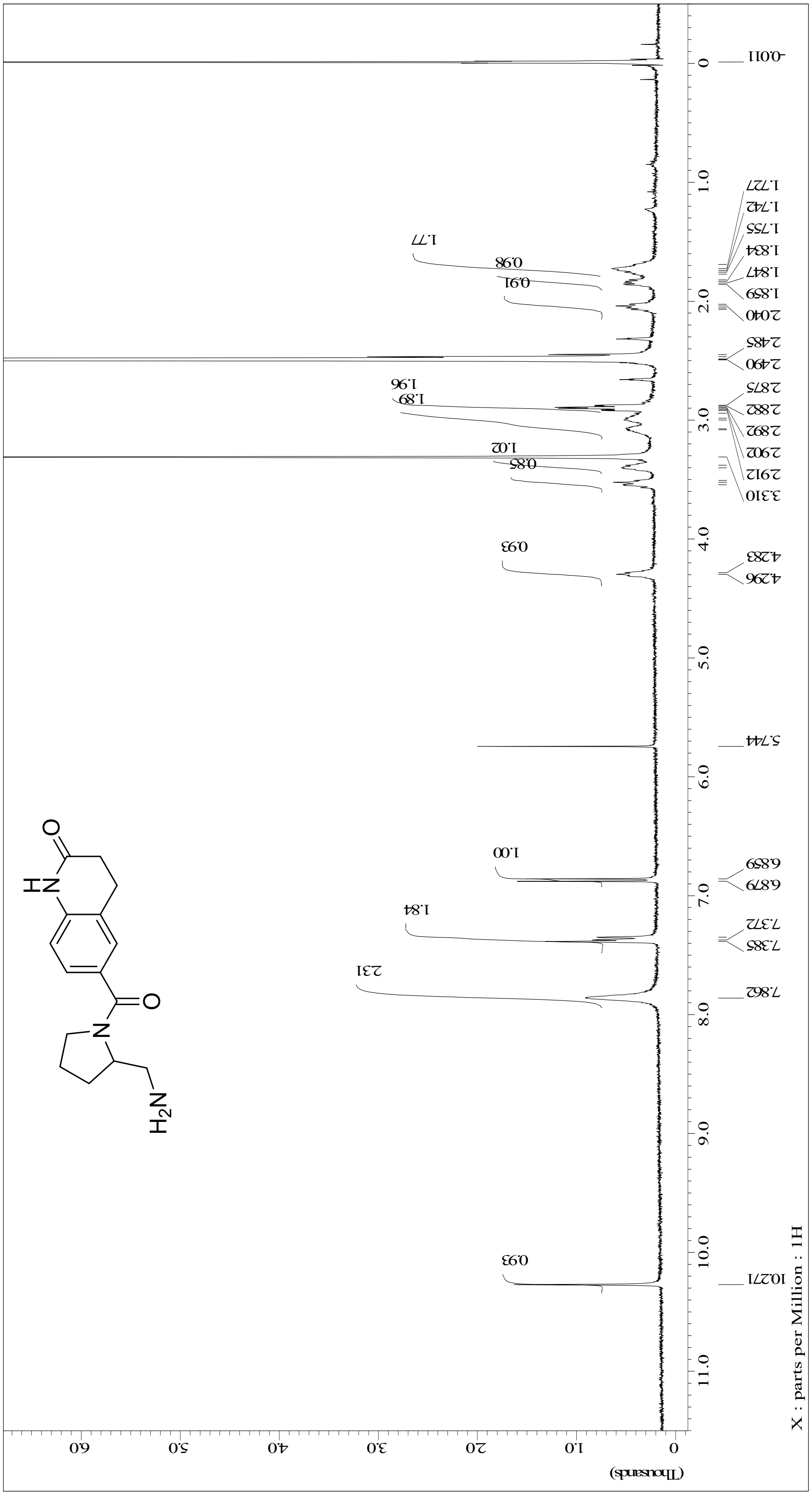
X : parts per Million : 1H
(Thousands)
10.271
7.862
7.385
7.372
6.879
6.859
5.744
4.296
4.283
3.310
2.912
2.902
2.892
2.882
2.875
2.490
2.485
2.040
1.859
1.847
1.834
1.755
1.742
1.727
-0.011
H$_2$N
NH
O

CS-e4027 in MeOD

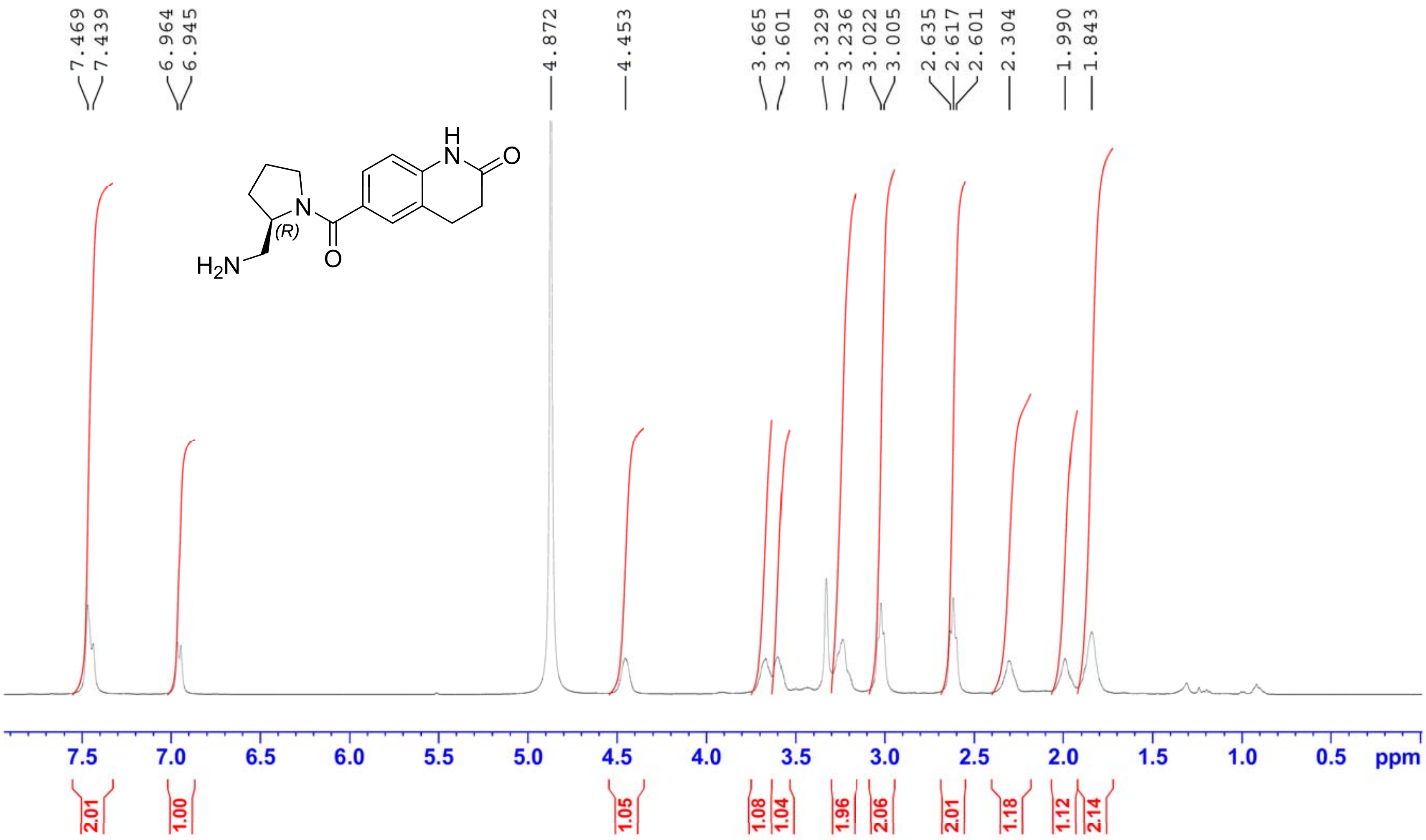

CS-e4025 in MeOD

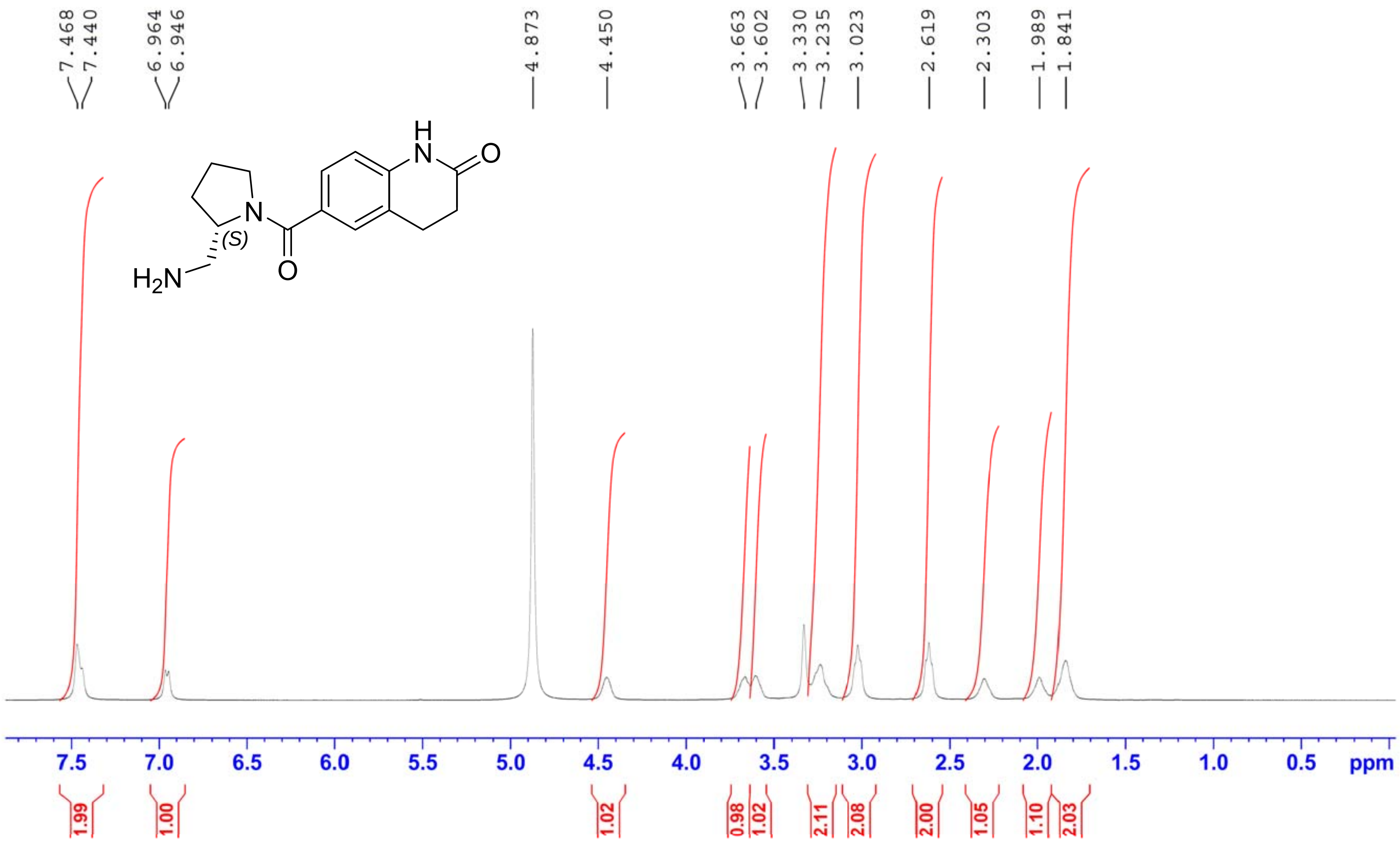

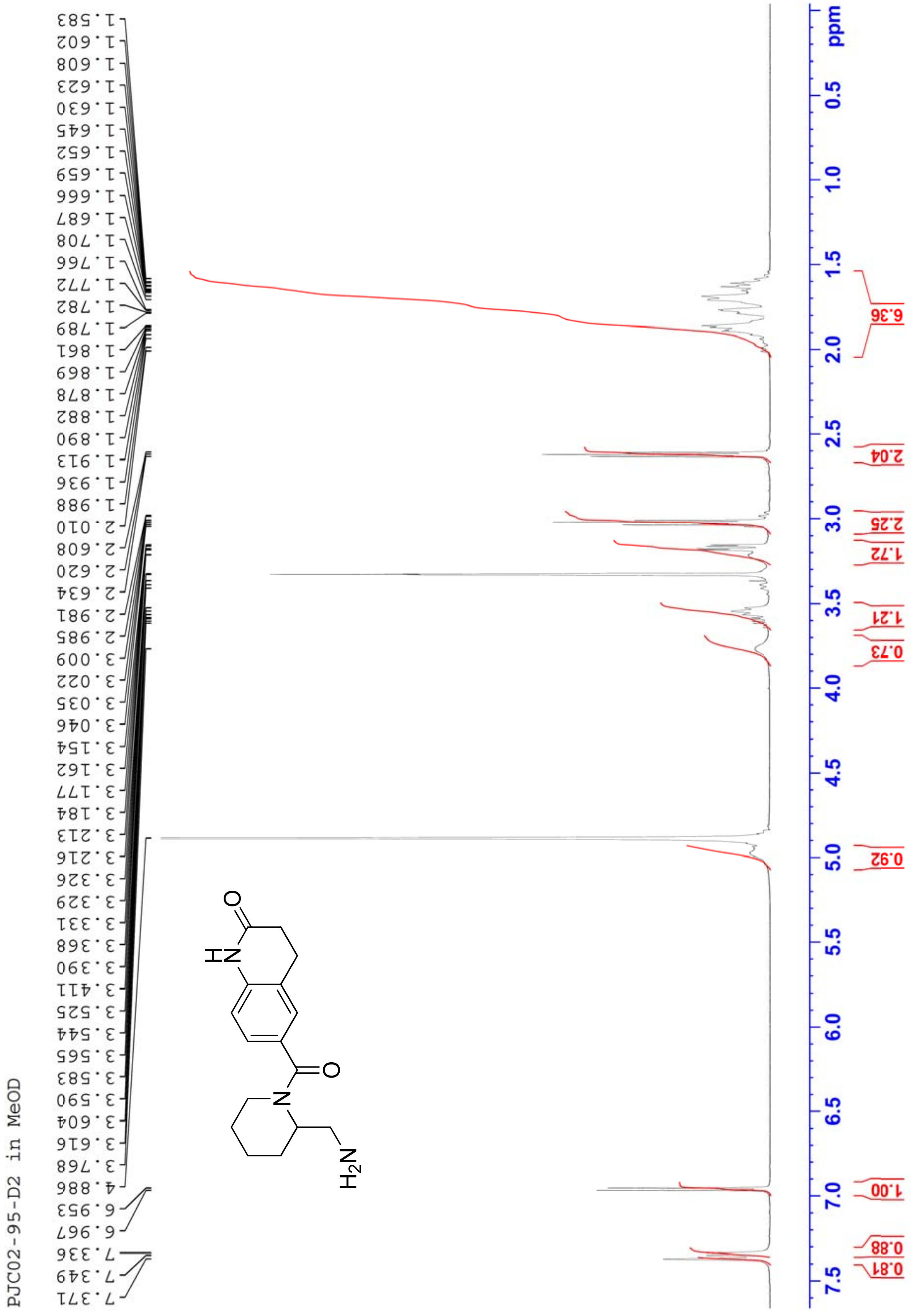
PJC02-95-D2 in MeOD
1.583
1.602
1.608
1.623
1.630
1.645
1.652
1.659
1.666
1.687
1.708
1.766
1.772
1.782
1.789
1.861
1.869
1.878
1.882
1.890
1.913
1.936
1.988
2.010
2.608
2.620
2.634
2.981
2.985
3.009
3.022
3.035
3.046
3.154
3.162
3.177
3.184
3.213
3.216
3.326
3.329
3.331
3.368
3.390
3.411
3.525
3.544
3.565
3.583
3.590
3.604
3.616
3.768
4.886
6.953
6.967
7.336
7.349
7.371
O
H
N
O
N
$H_2N$
6.36
2.04
2.25
1.72
1.21
0.73
0.92
1.00
0.88
0.81
ppm
0.5
1.0
1.5
2.0
2.5
3.0
3.5
4.0
4.5
5.0
5.5
6.0
6.5
7.0
7.5

PJC03-01-1 in MeOD

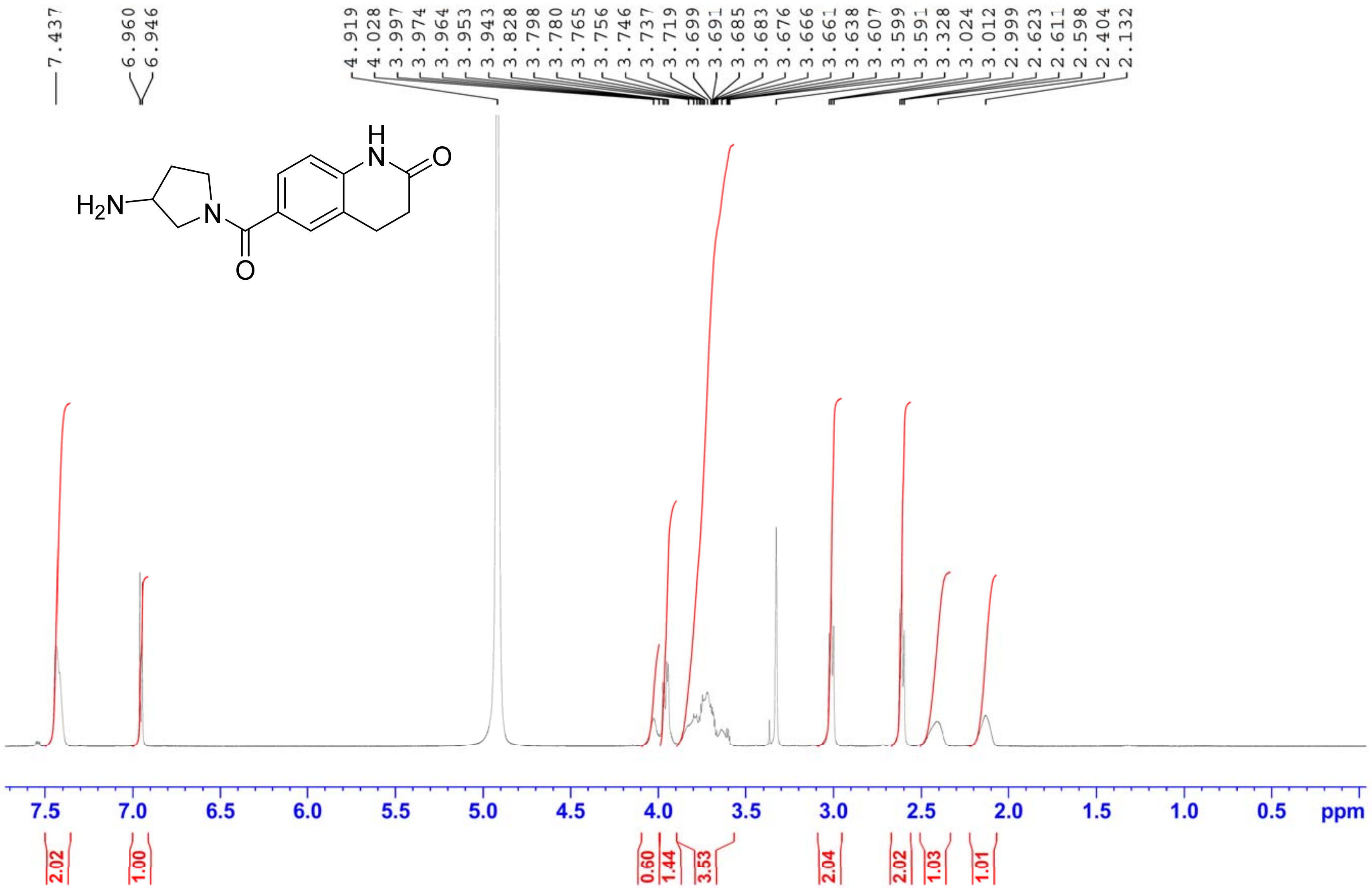

PJC03-02-1 in MeOD

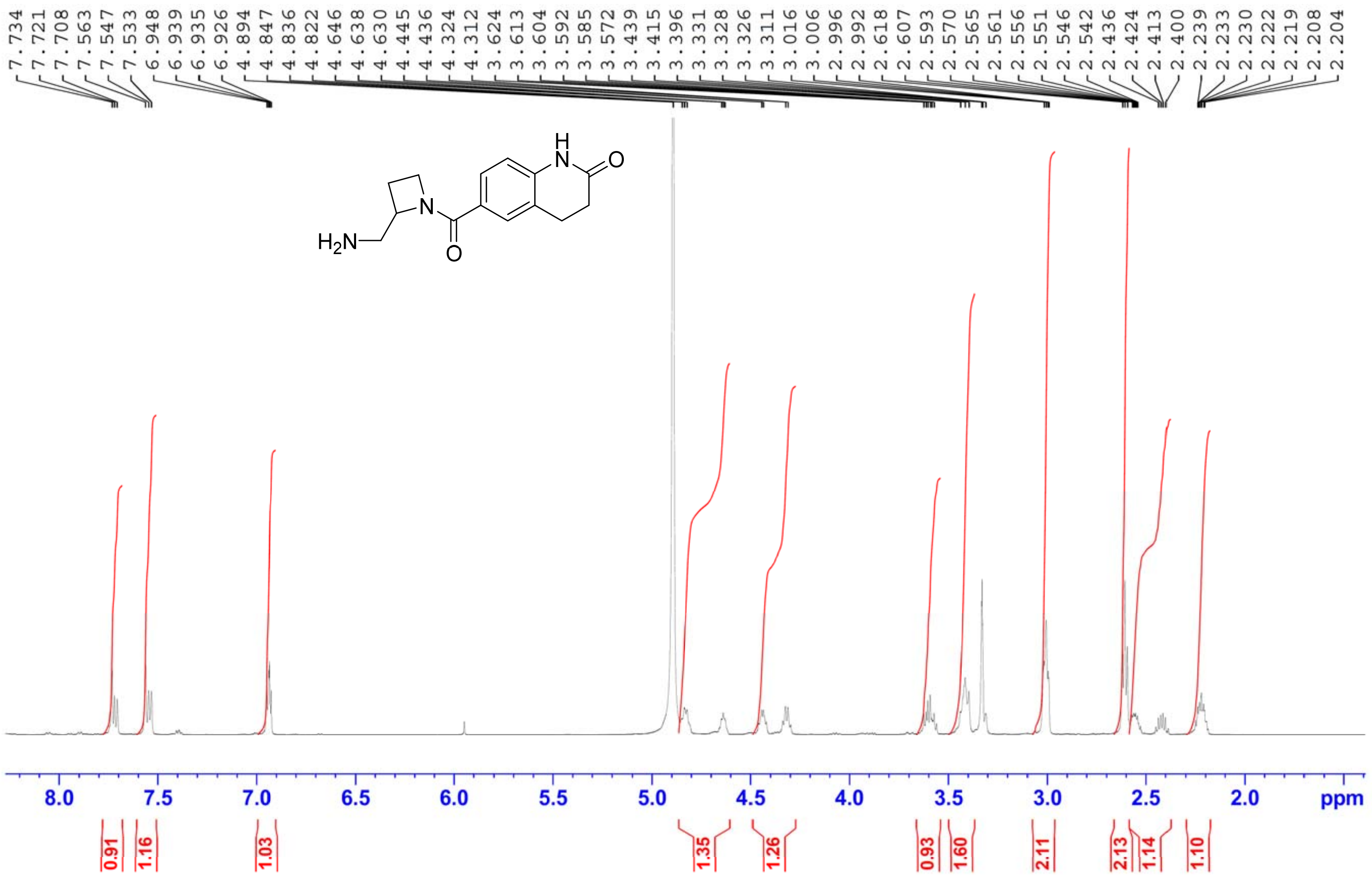

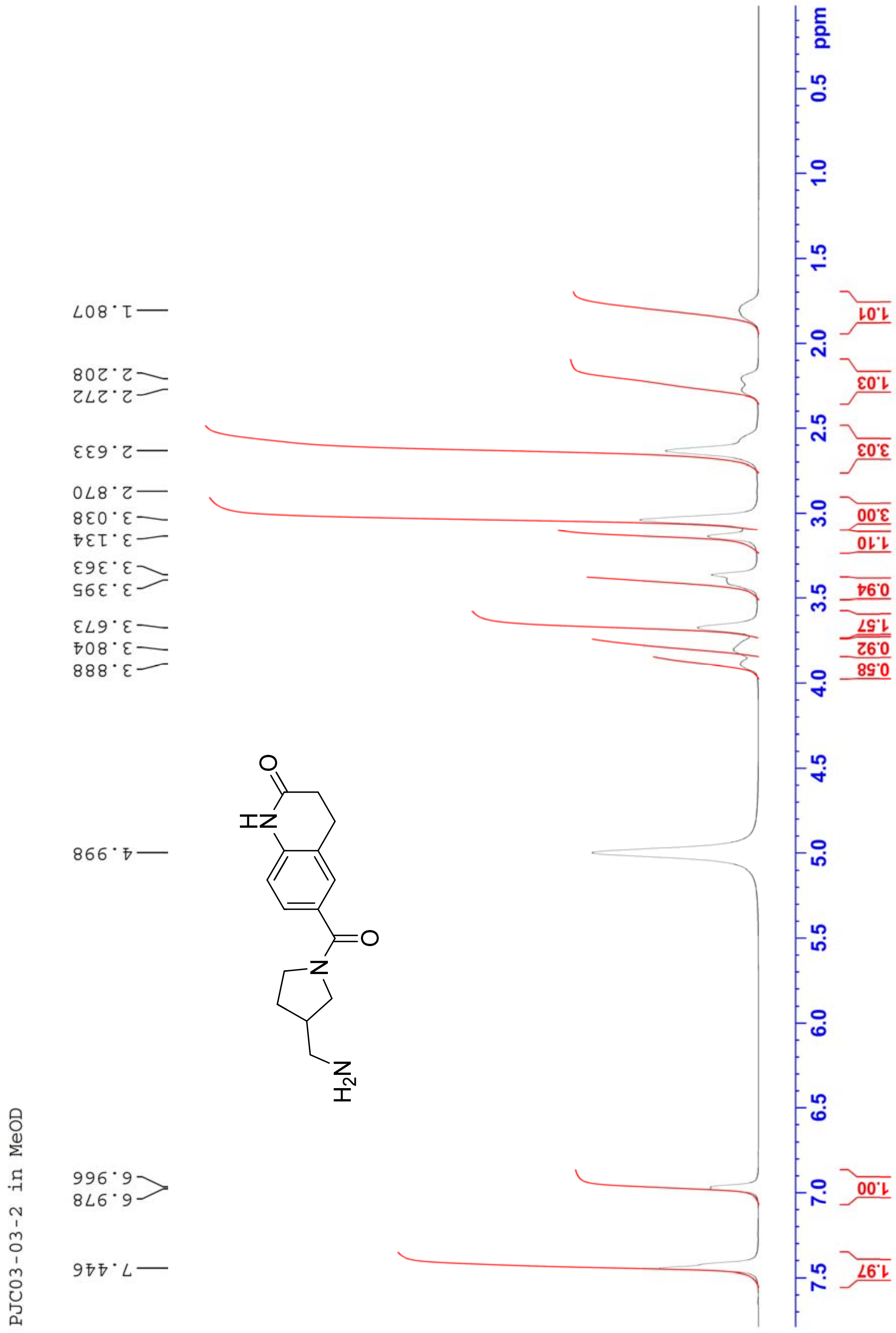
PJC03-03-2 in MeOD
7.446
6.978
6.966
4.998
3.888
3.804
3.673
3.395
3.363
3.134
3.038
2.870
2.633
2.272
2.208
1.807
NH
O
N
O
H2N
ppm
0.5
1.0
1.5
2.0
2.5
3.0
3.5
4.0
4.5
5.0
5.5
6.0
6.5
7.0
7.5
1.97
1.00
0.58
0.92
1.57
0.94
1.10
3.00
3.03
1.03
1.01

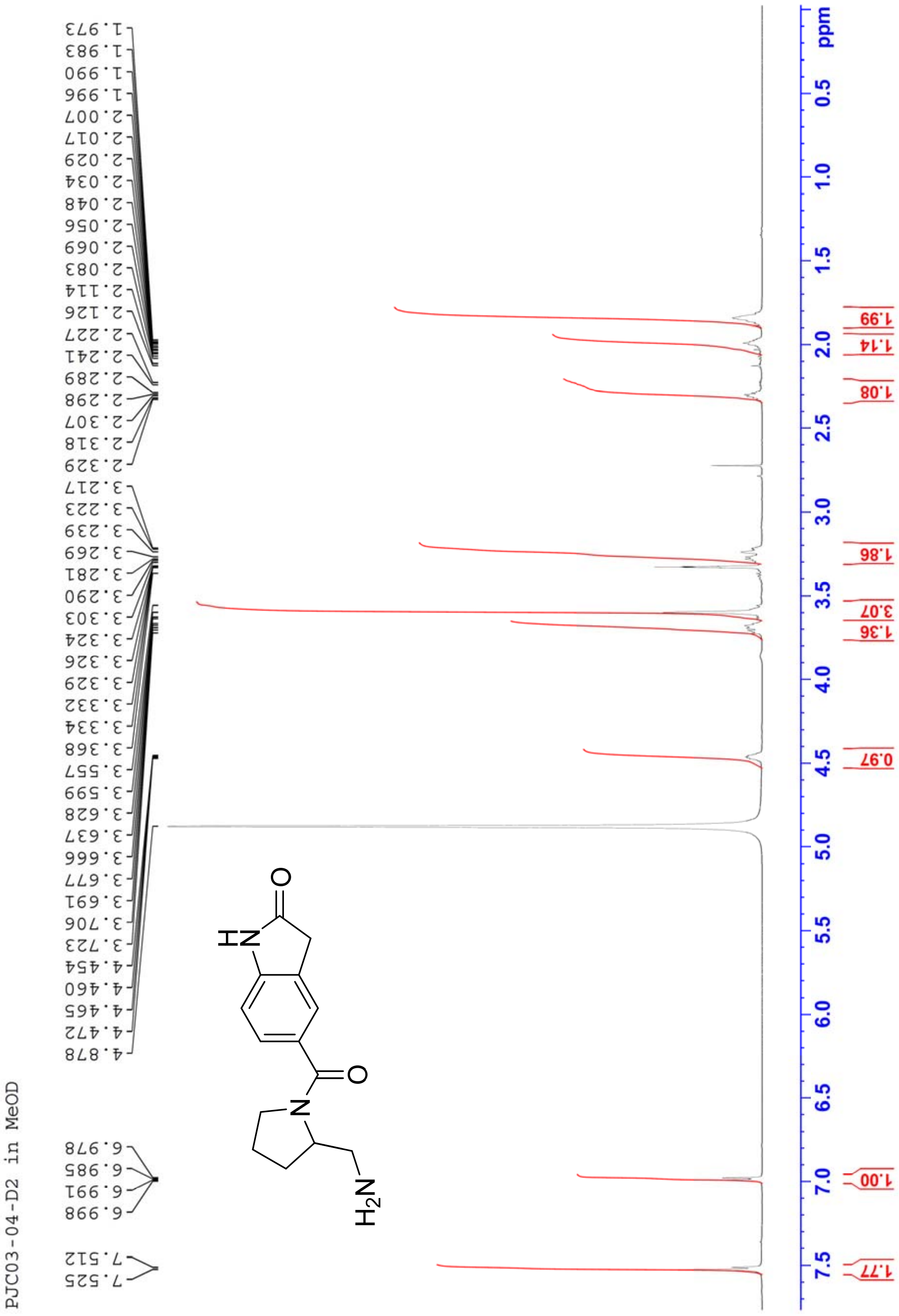
PJC03-04-D2 in MeOD
7.525
7.512
6.998
6.991
6.985
6.978
4.878
4.472
4.465
4.460
4.454
3.723
3.706
3.691
3.677
3.666
3.637
3.628
3.599
3.557
3.368
3.334
3.332
3.329
3.326
3.324
3.303
3.290
3.281
3.269
3.239
3.223
3.217
2.329
2.318
2.307
2.298
2.289
2.241
2.227
2.126
2.114
2.083
2.069
2.056
2.048
2.034
2.029
2.017
2.007
1.996
1.990
1.983
1.973
ppm
1.77
1.00
0.97
1.36
3.07
1.86
1.08
1.14
1.99

PJC03-18-3 in MeOD

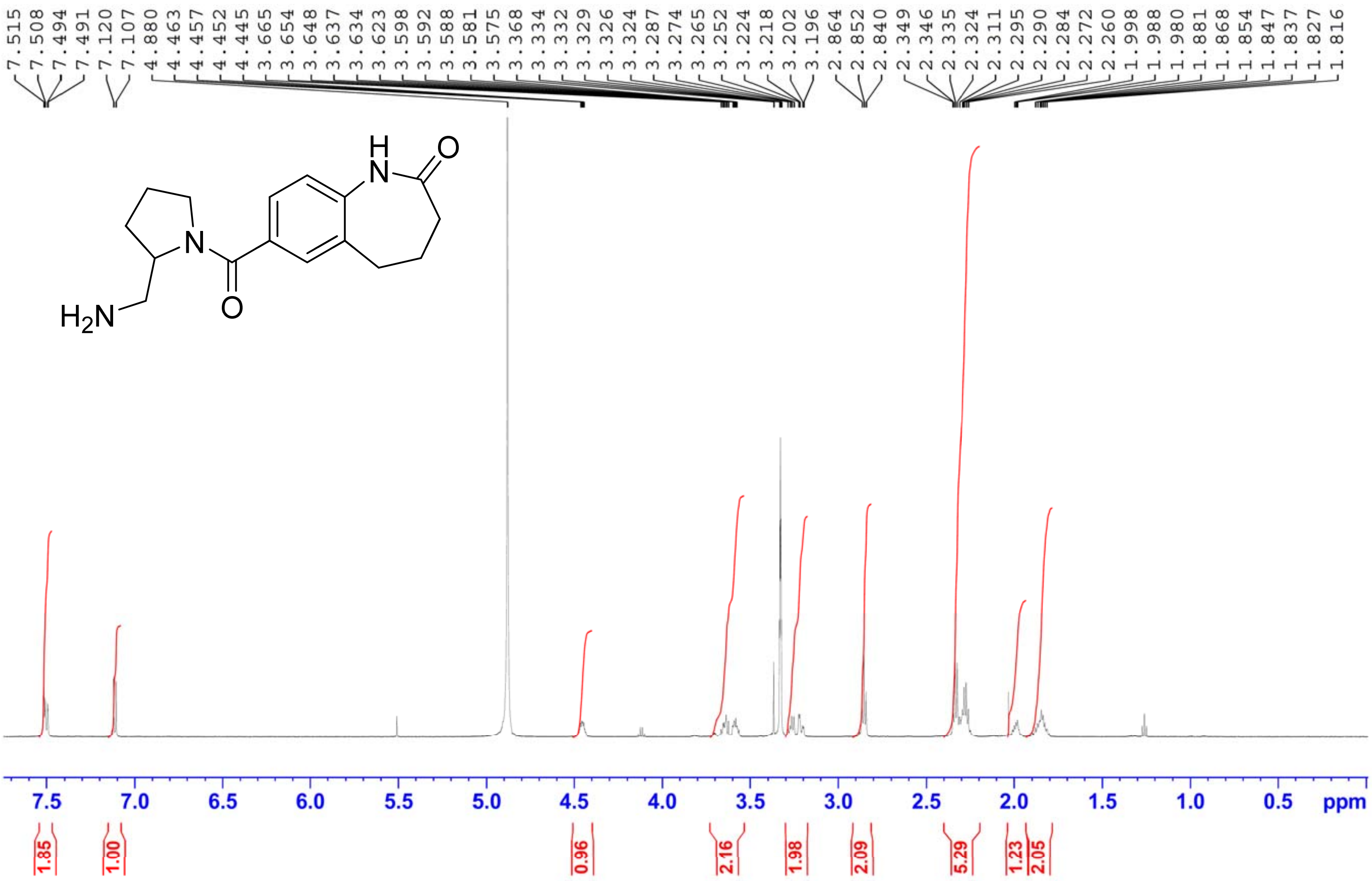

PJC03-05-1 A14 in MeOD

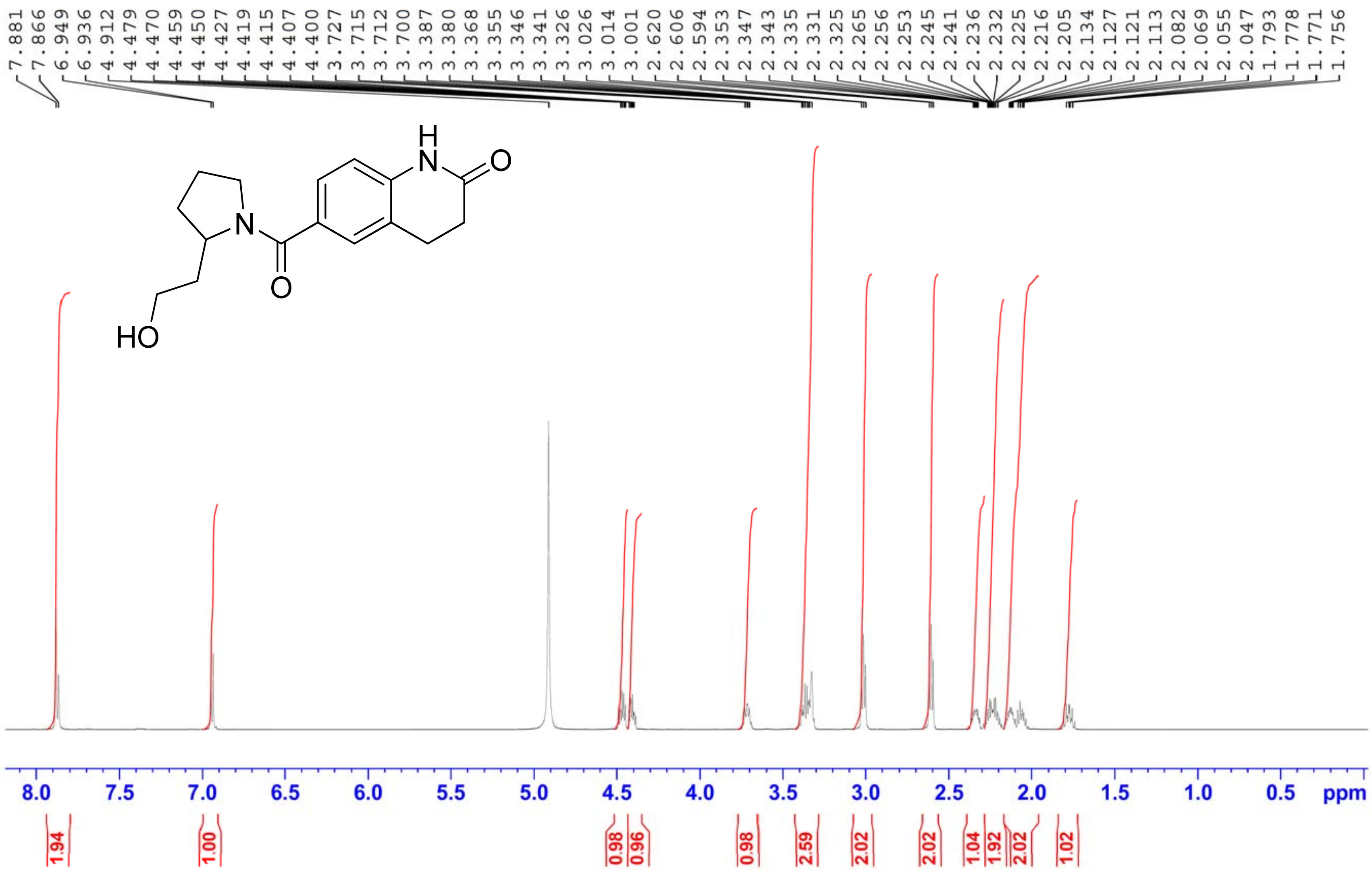

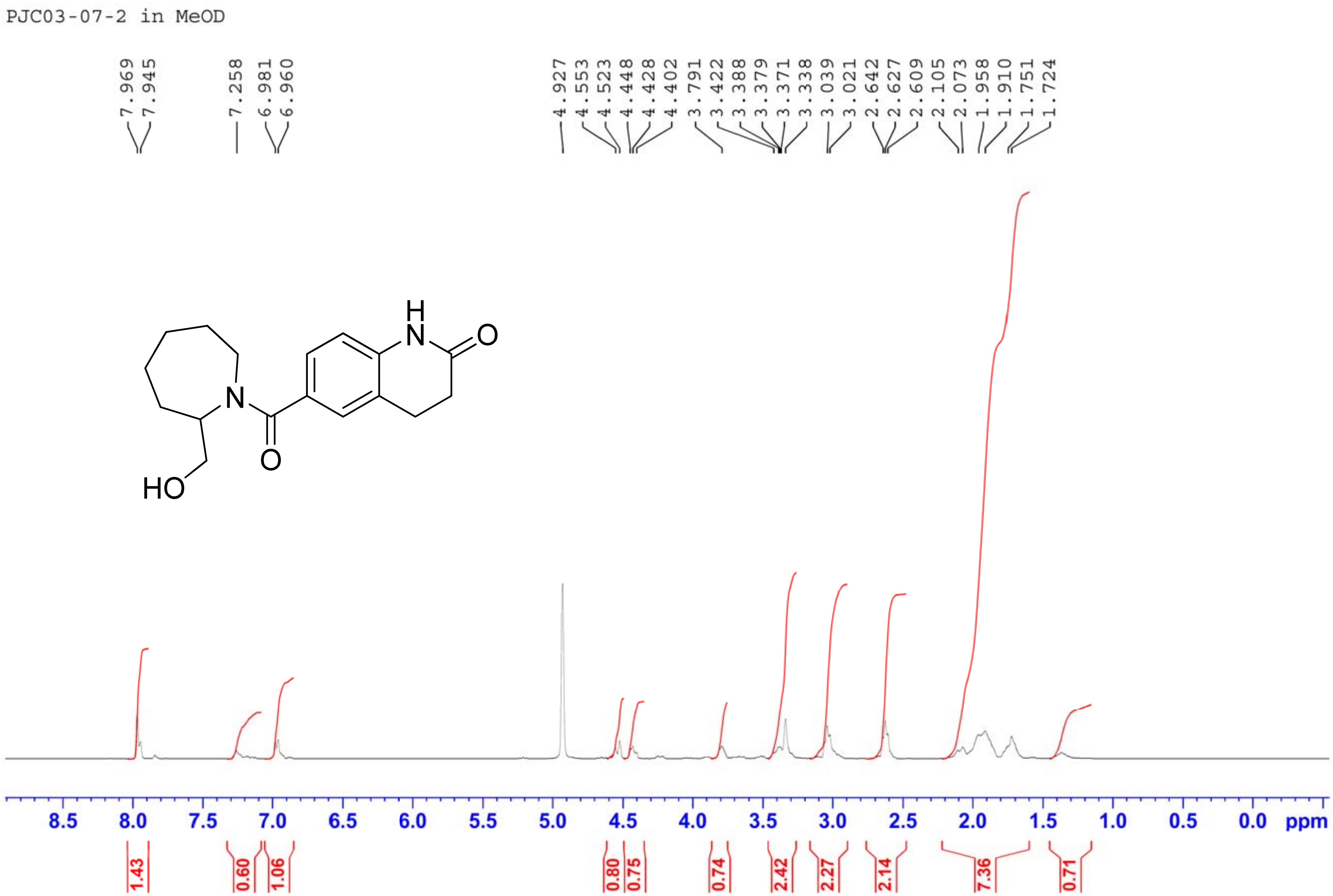
PJC03-07-2 in MeOD
7.969
7.945
7.258
6.981
6.960
4.927
4.553
4.523
4.448
4.428
4.402
3.791
3.422
3.388
3.379
3.371
3.338
3.039
3.021
2.642
2.627
2.609
2.105
2.073
1.958
1.910
1.751
1.724
H
N
O
N
O
HO
8.5
8.0
7.5
7.0
6.5
6.0
5.5
5.0
4.5
4.0
3.5
3.0
2.5
2.0
1.5
1.0
0.5
0.0
ppm
1.43
0.60
1.06
0.80
0.75
0.74
2.42
2.27
2.14
7.36
0.71

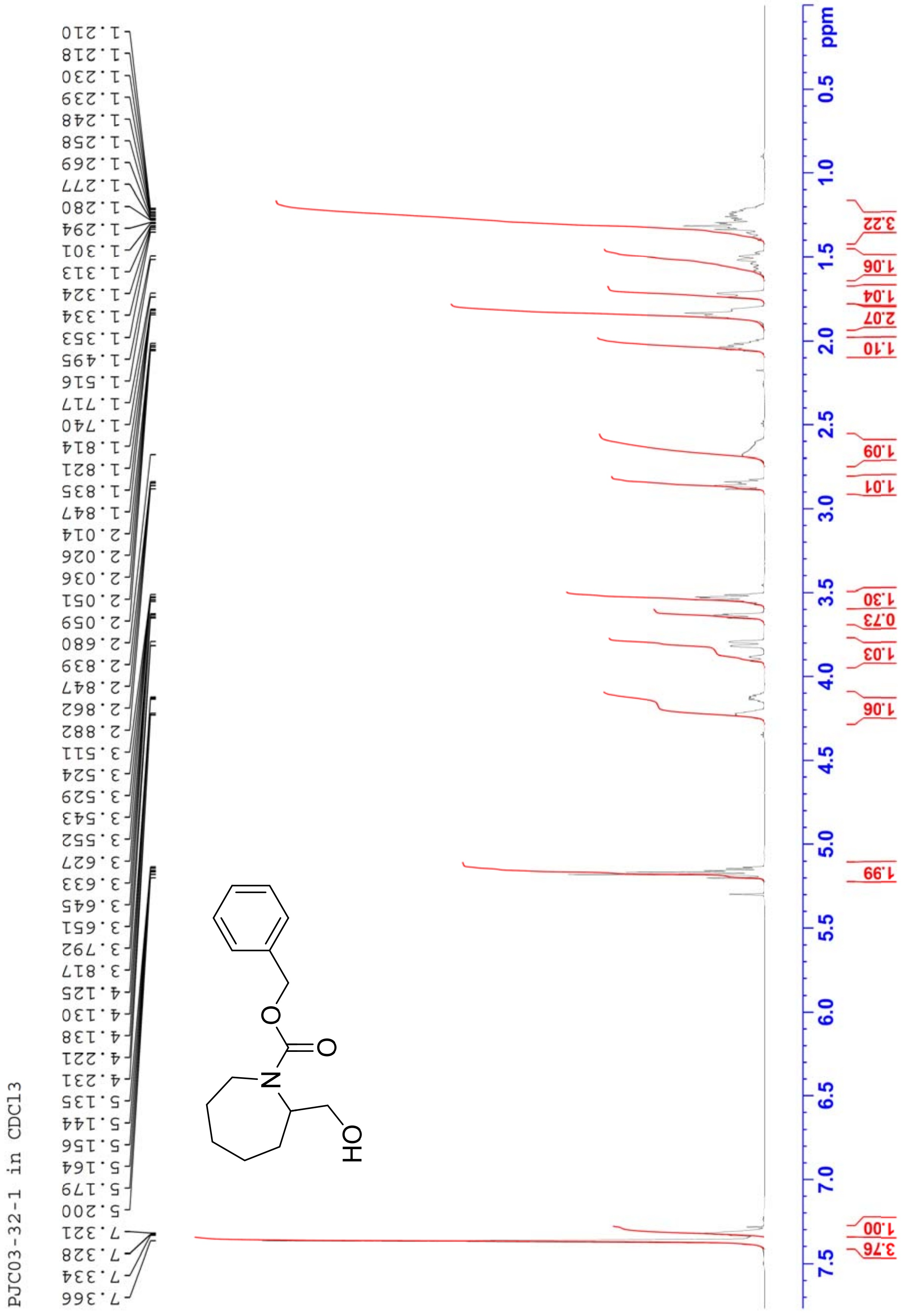
PJC03-32-1 in CDCl3
1.210
1.218
1.230
1.239
1.248
1.258
1.269
1.277
1.280
1.294
1.301
1.313
1.324
1.334
1.353
1.495
1.516
1.717
1.740
1.814
1.821
1.835
1.847
2.014
2.026
2.036
2.051
2.059
2.680
2.839
2.847
2.862
2.882
3.511
3.524
3.529
3.543
3.552
3.627
3.633
3.645
3.651
3.792
3.817
4.125
4.130
4.138
4.221
4.231
5.135
5.144
5.156
5.164
5.179
5.200
7.321
7.328
7.334
7.366
O
O
N
OH
ppm
0.5
1.0
1.5
2.0
2.5
3.0
3.5
4.0
4.5
5.0
5.5
6.0
6.5
7.0
7.5
3.22
1.06
1.04
2.07
1.10
1.09
1.01
1.30
0.73
1.03
1.06
1.99
1.00
3.76

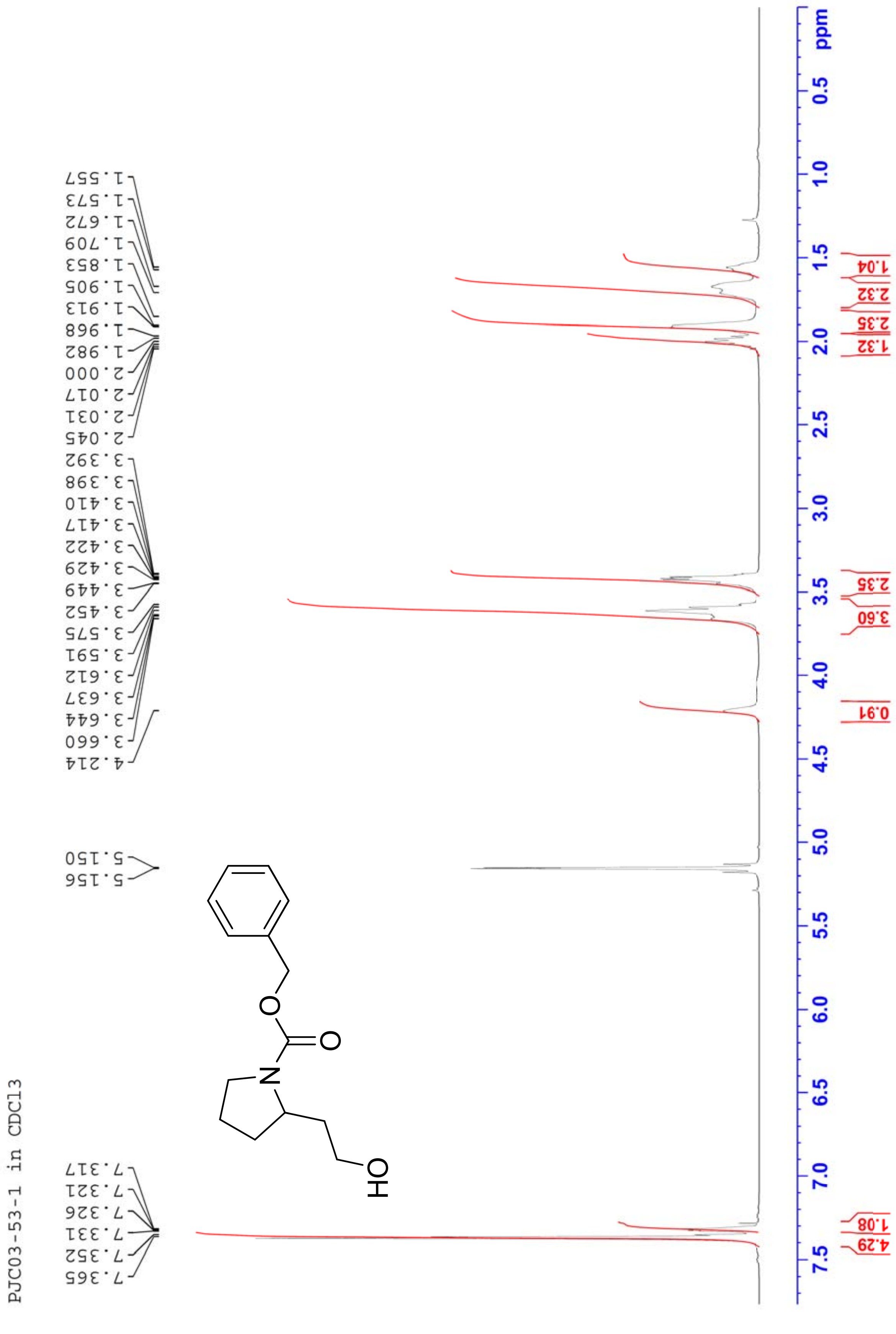
PJC03-53-1 in CDCl3
7.365
7.352
7.331
7.326
7.321
7.317
5.156
5.150
4.214
3.660
3.644
3.637
3.612
3.591
3.575
3.452
3.449
3.429
3.422
3.417
3.410
3.398
3.392
2.045
2.031
2.017
2.000
1.982
1.968
1.913
1.905
1.853
1.709
1.672
1.573
1.557
N
O
O
OH
ppm
4.29
1.08
0.91
3.60
2.35
1.32
2.35
2.32
1.04

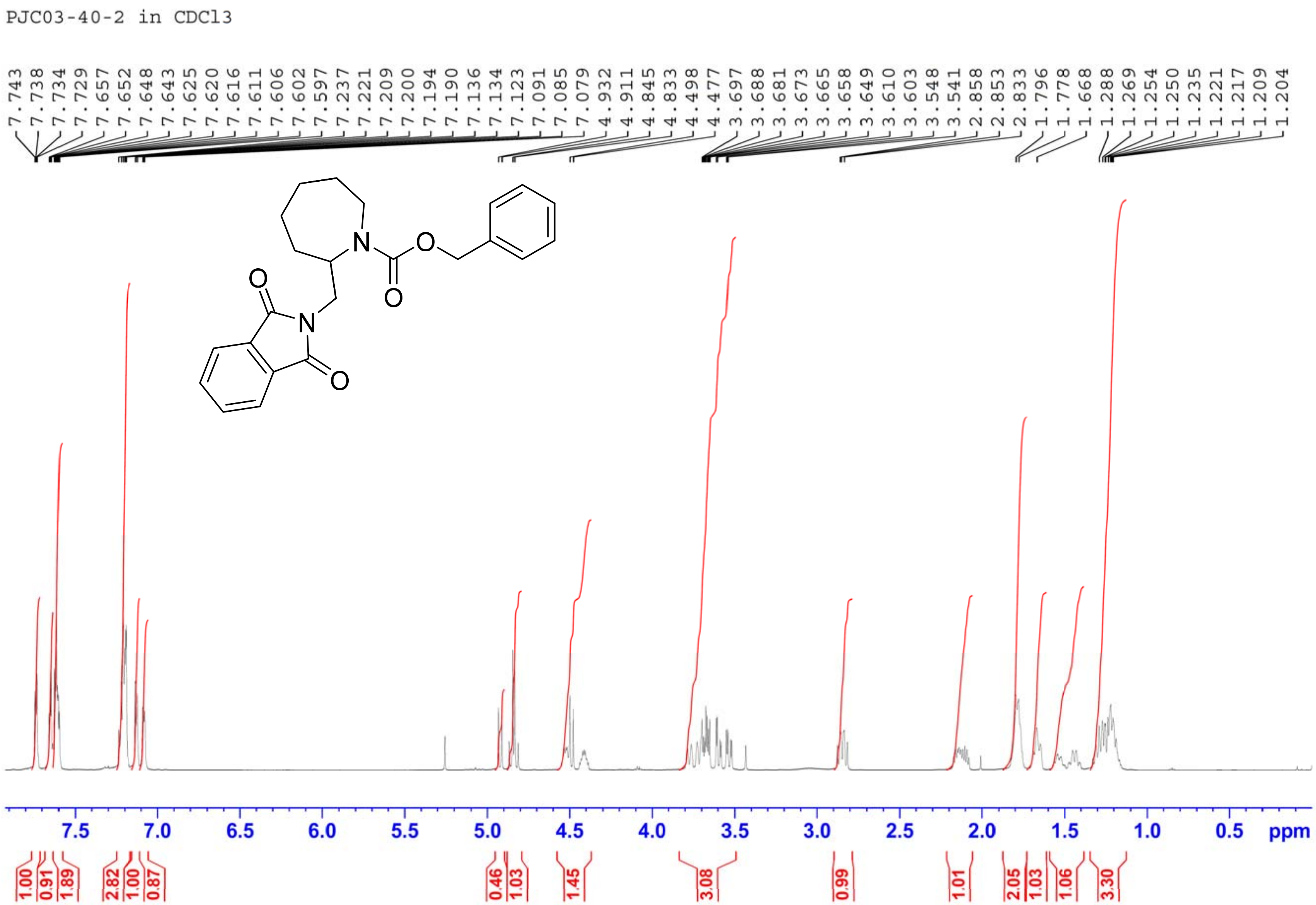
PJC03-40-2 in CDCl3
7.743 7.738 7.734 7.729 7.657 7.652 7.648 7.643 7.625 7.620 7.616 7.611 7.606 7.602 7.597 7.237 7.221 7.209 7.200 7.194 7.190 7.136 7.134 7.123 7.091 7.085 7.079 4.932 4.911 4.845 4.833 4.498 4.477 3.697 3.688 3.681 3.673 3.665 3.658 3.649 3.610 3.603 3.548 3.541 2.858 2.853 2.833 1.796 1.778 1.668 1.288 1.269 1.254 1.250 1.235 1.221 1.217 1.209 1.204
7.5 7.0 6.5 6.0 5.5 5.0 4.5 4.0 3.5 3.0 2.5 2.0 1.5 1.0 0.5 ppm
1.00 0.91 1.89 2.82 1.00 0.87 0.46 1.03 1.45 3.08 0.99 1.01 2.05 1.03 1.06 3.30

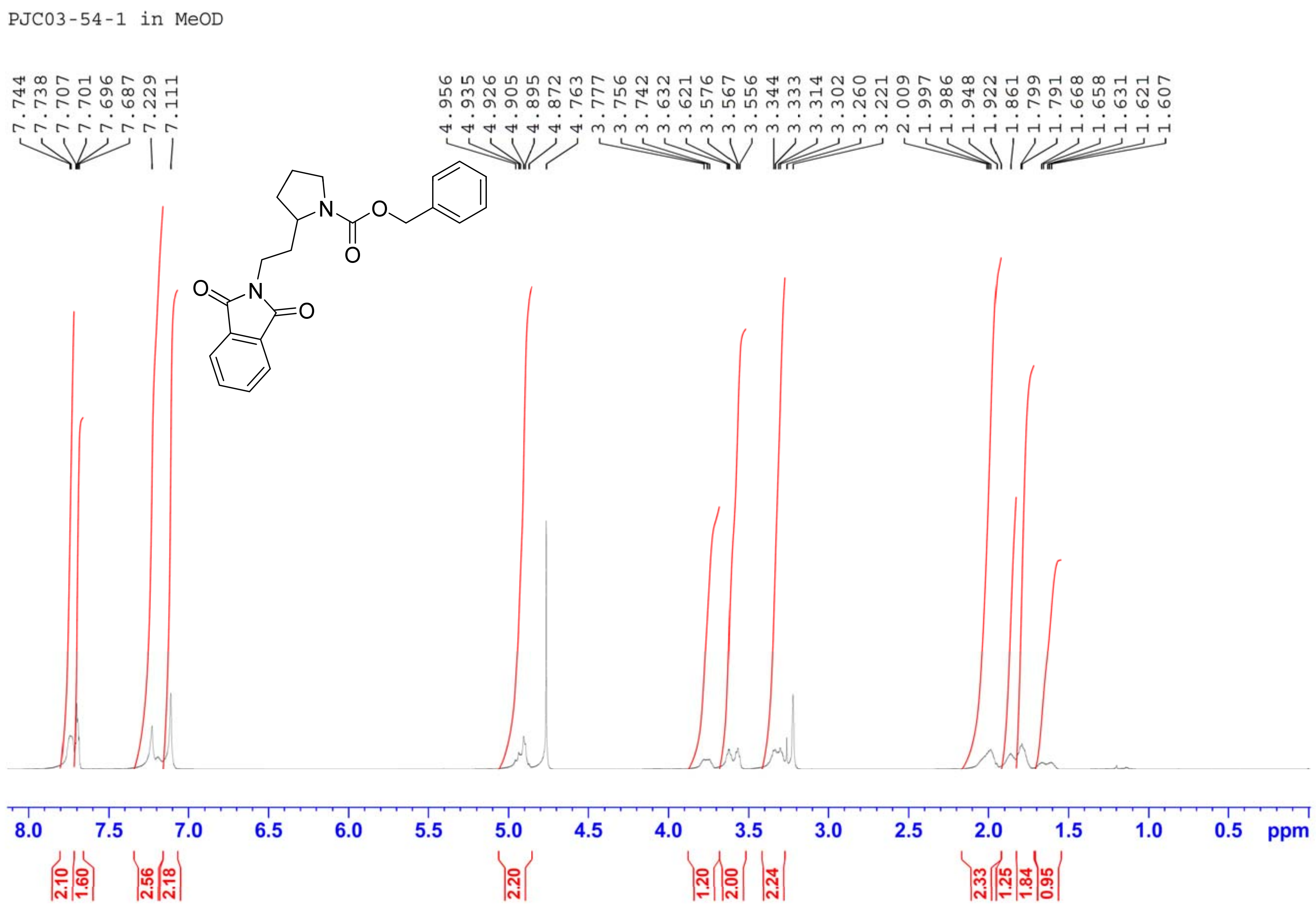
PJC03-54-1 in MeOD
7.744
7.738
7.707
7.701
7.696
7.687
7.229
7.111
4.956
4.935
4.926
4.905
4.895
4.872
4.763
3.777
3.756
3.742
3.632
3.621
3.576
3.567
3.556
3.344
3.333
3.314
3.302
3.260
3.221
2.009
1.997
1.986
1.948
1.922
1.861
1.799
1.791
1.668
1.658
1.631
1.621
1.607
N
O
O
N
O
O
8.0
7.5
7.0
6.5
6.0
5.5
5.0
4.5
4.0
3.5
3.0
2.5
2.0
1.5
1.0
0.5
ppm
2.10
1.60
2.56
2.18
2.20
1.20
2.00
2.24
2.33
1.25
1.84
0.95

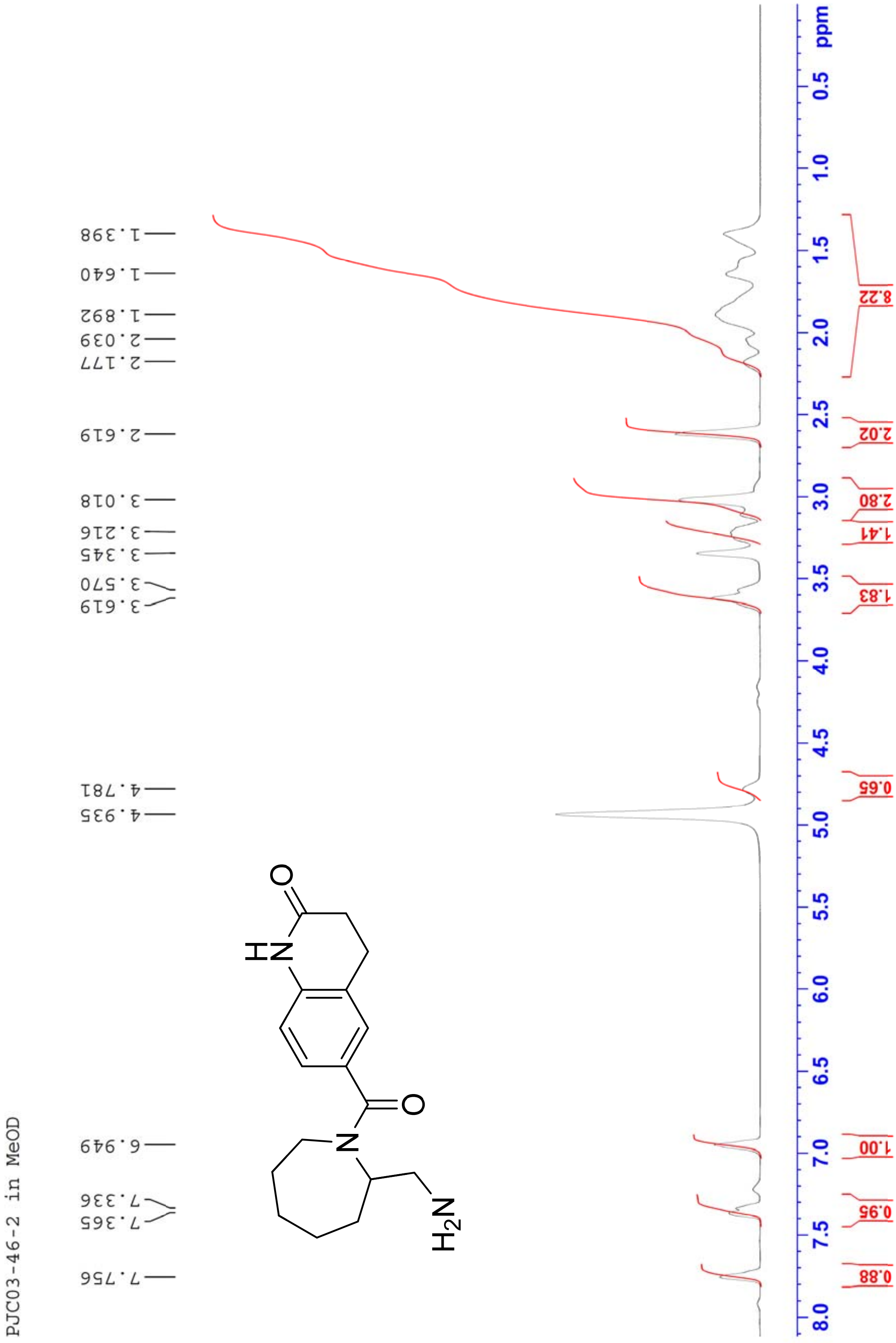

PJC03-46-2 in MeOD
7.756
7.365
7.336
6.949
4.935
4.781
3.619
3.570
3.345
3.216
3.018
2.619
2.177
2.039
1.892
1.640
1.398
NH
O
N
O
H2N
ppm
0.88
0.95
1.00
0.65
1.83
1.41
2.80
2.02
8.22
8.0 7.5 7.0 6.5 6.0 5.5 5.0 4.5 4.0 3.5 3.0 2.5 2.0 1.5 1.0 0.5

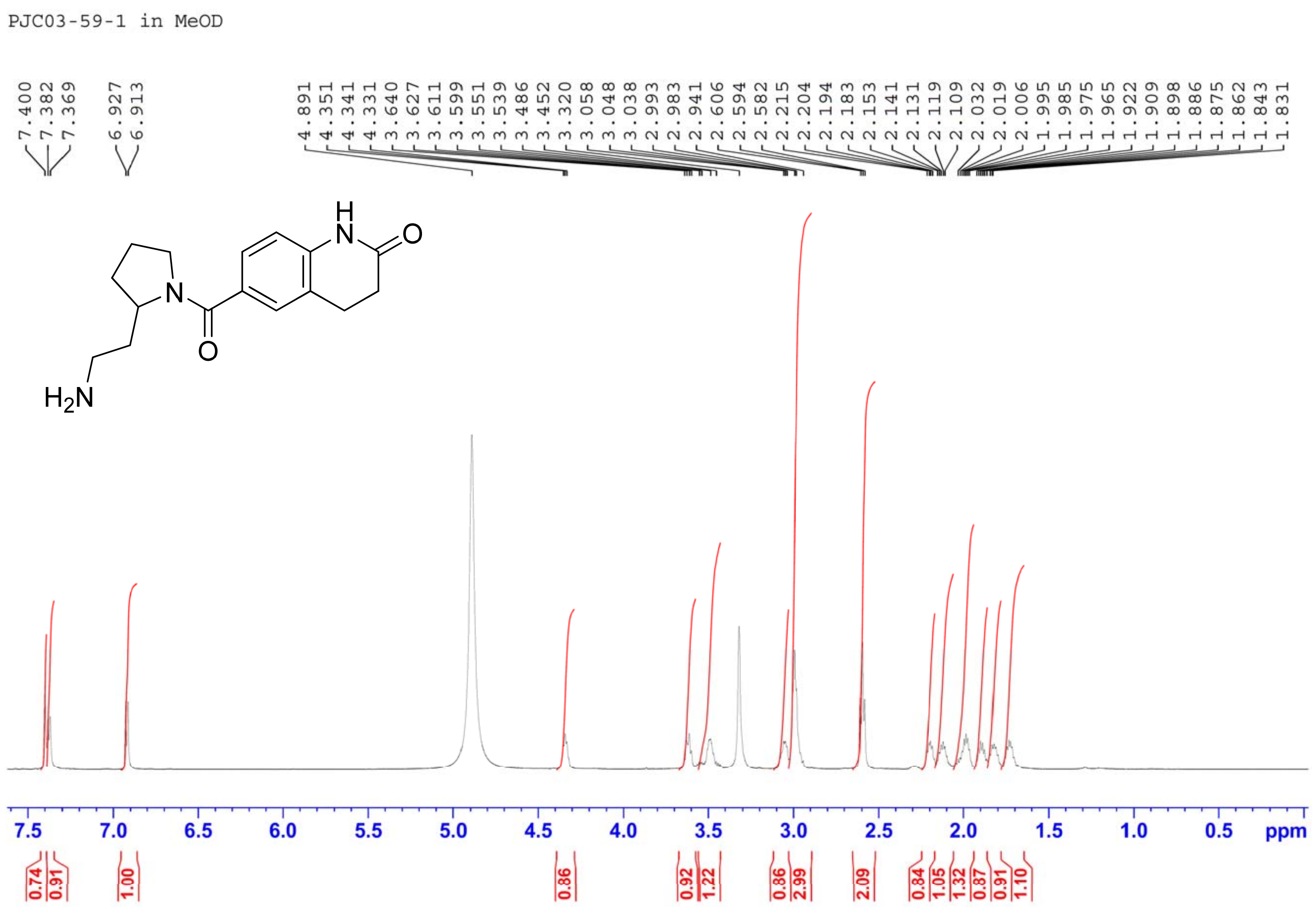
PJC03-59-1 in MeOD
7.400
7.382
7.369
6.927
6.913
4.891
4.351
4.341
4.331
3.640
3.627
3.611
3.599
3.551
3.539
3.486
3.452
3.320
3.058
3.048
3.038
2.993
2.983
2.941
2.606
2.594
2.582
2.215
2.204
2.194
2.183
2.153
2.141
2.131
2.119
2.109
2.032
2.019
2.006
1.995
1.985
1.975
1.965
1.922
1.909
1.898
1.886
1.875
1.862
1.843
1.831
H
N
O
N
O
$H_2N$
7.5
7.0
6.5
6.0
5.5
5.0
4.5
4.0
3.5
3.0
2.5
2.0
1.5
1.0
0.5
ppm
0.74
0.91
1.00
0.86
0.92
1.22
0.86
2.99
2.09
0.84
1.05
1.32
0.87
0.91
1.10
